# Annelid brain and nerve cord in siboglinid *Riftia pachyptila*

**DOI:** 10.1101/325431

**Authors:** Nadezhda N. Rimskaya-Korsakova, Sergey V. Galkin, Vladimir V. Malakhov

## Abstract

Vestimentifera is a peculiar group of marine gutless siboglinids which has uncertain position in annelid tree. The detailed study of the fragmentary explored central nervous system of vestimentiferans and other siboglinids is requested to trace the evolution of the siboglinid group. Among all siboglinids the vestimentiferans preserve the gut rudiment what makes them a key group to homologize main cerebral structures with the ones of typical annelids, such as supra- and subesophageal commissures, cirsumesophageal connectives etc. Histologically we revealed main annelid brain structures in the compact large brain of *Riftia pachyptila*: circumesophageal connectives (longitudinal nerve tracts) and commissures (dorsal, supra- and subenteral commissures). Innervation of tentacles makes them homologous to peristomial palps of the rest annelids. The single nerve cord is represented by paired intraepidermal longitudinal strands associated with the ventral ciliary field in vestimentum and bearing giant axons originating from at least four pairs of perikarya. The absence of regularly positioned ganglia and lateral nerves in the nerve cord in vestimentum and trunk and presence of them in the opisthosome segments. Among siboglinids, the vestimentiferans distinguished by a large and significatly differentiated brain which is reflection of the high development of the palp apparatus. *Osedax*, frenulates and *Sclerolinum* have less developped brain. Frenulates and *Sclerolinum* have good ganglionization in the opisthosome, which probably indicates its high mobility. Comparative neuroanatomical analysis of the siboglinids and annelid sister clades allows us to hypothesize that the last common ancestor of siboglinids might had brain with a dorsal commissure giving rise neurite bundles to palps and paired ventral nerve cord.

## Introduction

Vestimentifera is a peculiar group of marine gutless annelids inhabiting mainly areas of hydrothermal vents and hydrocarbon seeps [1–3]. The first anatomical details of the nervous system of Vestimentifera was made in the description of the first discovered vestimentiferan *Lamellibrachia barhami* [4]. Later on, nervous system was studied in *L. luymesi* [5,6], *Riftia pachyptila* [7–9], *Ridgeia piscesae* [10], *Oasisia alvinae* [11], *L. satsuma* [12]. By means of light microscopy and histology it was shown the presence of the ventral nerve cords and positions of perikariya and neuropile in brain of larval [13,14] and adult vestimentiferans [6,9–12,15]. Electron microscopical studies revealed presence of the sensory cells and glial cells structuring neuropile, and form a myelin sheath around the giant axons [9]. In adult vestimentiferans the brain occupies an unusual for annelids antero-ventral position. Jones and Gardiner [13] found that in juveniles the rudiment of the brain is laid in the base on the dorsal side of the oral siphon. After the reduction of the oral siphon, the brain rudiment shifts to the antero-ventral position. Jones and Gardiner [8] suggested that the brain of vestimentiferans is a result of the fusion of the supra- and subesophageal ganglia and the circumesophageal connectives. Based on the fact that a coelomic channel passes through the brain in which the rudimentary intestine remains in young individuals, this assumption seems very likely. However, it needs to be confirmed by a more detailed comparison of the intracerebral structures of vestimentiferans and more typical annelids.

Vestimentiferan tubeworms together with Frenulata [4], *Sclerolinum* [16] and *Osedax* [17] refer to annelid group Siboglinidae [18]. The nervous system of the vestimentiferans and the rest siboglinids was studied by various methods at the different levels of detalization what makes them difficult to compare. The architecture of frenulates’ central nervous system is known based on histological and histochemical studies of the ventral nerve cords and rings and brain area of early branched species *Siboglinum caulleryi*, *S. fiordicum* and *Nereilinum murmanicum, and* derived ones as *Polybrachia annulata, Spirobrachia grandis* [18–24]. Electron microscopy revealed presence of glial and sensory elements in epidermis of frenulates [25]. Structure of central nervous system of females and dwarf males of *Osedax* is described by means of immunohistochemistry combined with confocal microscopy that revealed numerous commissures and connectives in the brain and trunk nervous system [26,27]. Semi-thin sectioning and light microscopy revealed in brain of *Sclerolinum contortum* the layers of apical perikarya and basal neuropile [28]. Precise ultrastructural neural studies on *Osedax* and *Sclerolinum* were not yet made. Thus, the degree of anatomical study of the organization of the nervous system of vestimentiferans and other siboglinids remains fragmentary and insufficient to make meaningful comparisons with the nervous system of annelids, to which siboglinids are close, according to phylogenetic data [29–34]. Detailed neural reconstructions by comparable methods of siboglinids are higly requested to identify key features and trace their neural evolution.

Phylogenetic position of Vestimentifera and the whole group Siboglinidae in the annelid system remains controversial. Various annelid sister groups have been proposed, e.g. Oweniidae [35,36], Sabellidae [31,37], Cirratuliformia [38,39], or Clitellata [40,41]. Nervous system of the listed annelids is described by various authors [42–52], but there is still no attempts to reveal if there is any neural similarities among siboglinids and proposed sister groups of annelids.

Among all siboglinids the juvenile of vestimentiferans preserve the gut rudiment, so it is the key group to homologize the supra- and subesophageal brain parts of ventral brain of siboglinids with the typical annelid brains. Moreover, vestimentiferans are distinguished by their enormous sizes (*Riftia* reaches 1,5 m in length) what makes their histological studies very informative for 3D reconstructions. The main goal of the current work is to reconstruct organization of the central nervous system of vestimentiferan tubeworm *Riftia pachyptila* with special accent to its brain structure. This data is necessary for comparison of neuroanatomy of vestimentiferan tubeworms and sister groups of annelids to find the possible ancestor features in the nervous system of siboglinids.

## Materials and Methods

### Collection and Fixation

Five specimens of *Riftia pachyptila* Jones, 1981 [7] were collected at different latitudes of the East Pacific Rise (EPR), including the Guaymas Basin, Gulf of California, by the *Pisces* manned submersible during the 12th cruise of RV Akademik Mstislav Keldysh in 1986 and by *Mir*-1 & 2 manned submersibles during its 49th cruises in 2003. Lengths of examined specimens are from 8 to 808 mm. In Table 1 there is the data on collection sites and sexes of specimens.

**Table 1.** The studied specimens collected during cruises of the RV *Akademik Mstislav Keldysh* (AMK).

| specimens |  |  | collection sites |  |  |
| --- | --- | --- | --- | --- | --- |
| # | sex | length, mm | name & coordinates | depth, m | # station, ROV, year |
| 1 | juvenile | 8 | 9°N EPR:<br>09° 50,53' N, 104°17,51' W | 2552 | AMK-4668, *Mir-1*, 2003 |
| 2 | female | 16 | Guaymas Basin:<br>27° 02,45' N, 111°22,80' W | 1990 | AMK-1519, *Pisces-VII*, 1986 |
| 3 | female | 34 | 9°N EPR:<br>09° 50,52' N, 104°17,52' W | 2524 | AMK - 4623, *Mir-1*, 2003 |
| 4 | male | 79 | 9°N EPR:<br>09° 50,52' N, 104°17,52' W | 2524 | AMK-4623, *Mir-1*, 2003 |
| 5 | female | 808 | Guaymas Basin:<br>27° 00,47' N, 111°24,57' W | 2001 | AMK-4714, *Mir-2*, 2003 |

### Histology & LM photography

Four animals used for anatomical analysis were fixed in Bouin’s solution and stored in 70% ethanol. The material was processed by the standard histological procedure, including dehydration in alcohols and embedding in paraffin, paraplast, or histowax. Transverse sections (5 and 7 μm) were produced with a Leica RM 2125 microtome (Leica Microsystems, Wetzlar, Germany), stained with Caracci hematoxylin, and examined under a Zeiss Axioplan2 microscope equipped with AxioCam HRm camera (Carl Zeiss Microscopy, LLC, United States) as well as Leica DM5000 B equipped with Leica DFC425 C camera. Microscopic images optimized for contrast and level in Adobe Photoshop 7.0 (Adobe Systems, San Jose, CA, USA). Drawings were performed with Adobe Illustrator CC 2014. For visualization of the anastomosing neurites in the trunk epidermis a specimen of 808 mm was pictured by Canon Power Shot S90 camera.

### 3D modeling

Arrangements of neurite bundles in the brain and the anteriormost ventral nerve cord were visualized with the software 3D-DOCTOR 3.5.040724 (Able Software Corporation of Lexington, USA). Alignment was performed in the same software with comparing the sections of adjacent planes. Image seria of 77 cross sections of the 78 mm specimen was used for modeling of the brain organization. 19 objects were traced inside brain, including boundary of the brain. Photos are saved in JPEG format with a resolution of 3900 × 3090 pixels and 8 bits / pixel. The field of view is 2812.00 μm, the parameters of the voxels of the images are 0.721026 × 0.721026 × 15 μm3. On the basis of the outlined boundaries three-dimensional models were obtained. The smoothing tool used for natural perception of the surface of objects. Interactive features as well as transparency filter, different colours and lighting effects applied to show complex and hidden objects. Three-dimensional images under appropriate angles were processed in Adobe Photoshop 7.0 (Adobe Systems, San Jose, CA, USA).

## Results

### Gross anatomy of nervous system

*Riftia’s* central nervous system is composed of a ventral **brain** and **ventral nerve cord** (B, *VNC*, Figs 1A, 2A, 3A, 4A).

**Fig 1.**
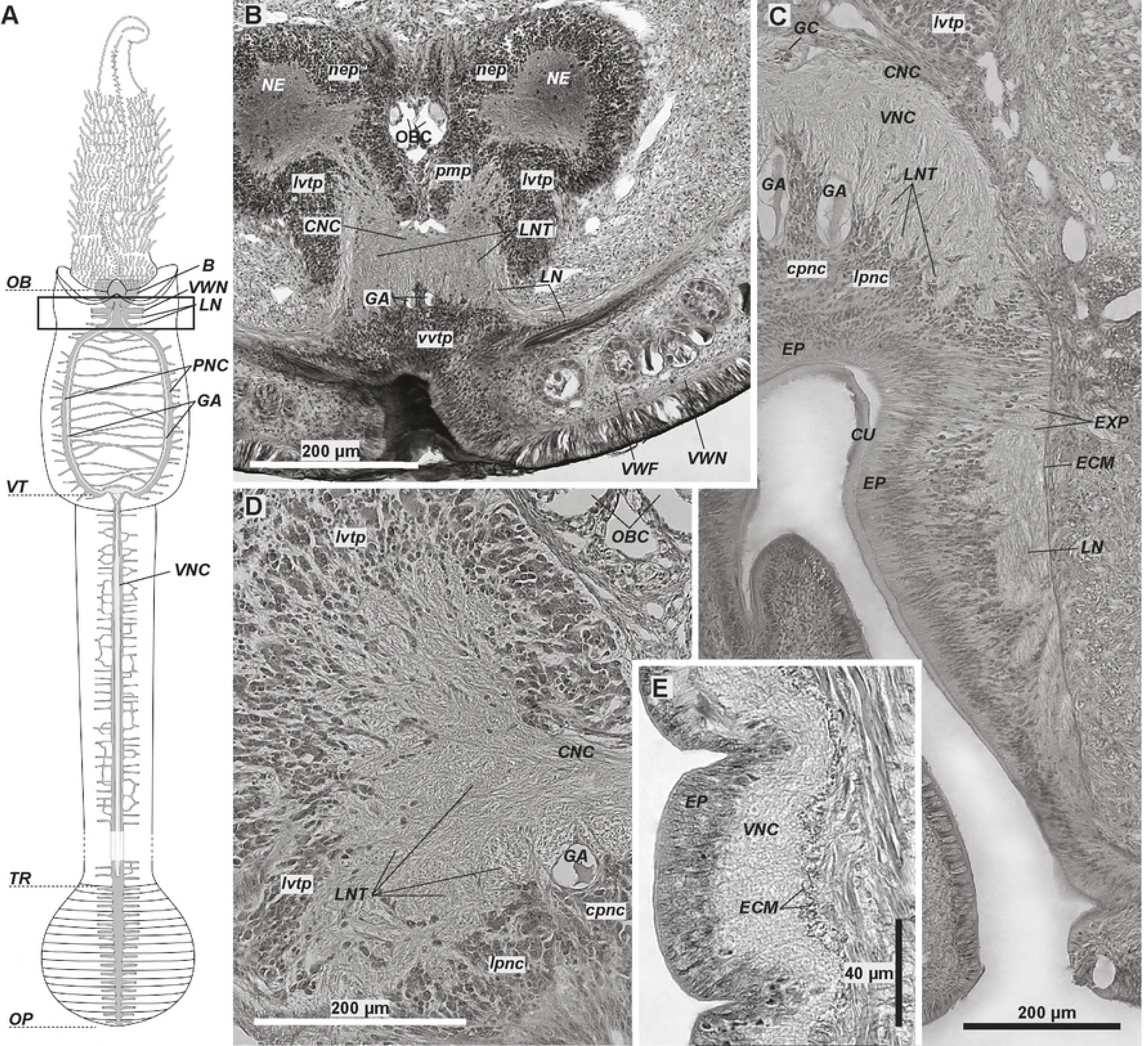
Anterior part of the ventral nerve cord of *Riftia pachyptila*. A - scheme of the central nervous system which main elements are in grey, and giant axons are in light grey. The frame indicates the area corresponding to histological cross (B-D) and parasagittal (E) sections. Dotted lines show the region borders. B - posteriormost brain; elements of the ventral nerve cord projecting into the brain. C - ventral nerve cord (*VNC*) just posterior to the brain. D - longitudinal nerves in the transition of the ventral nerve cord and brain. E - the intraepidermal ventral nerve cord (*VNC*). *B* – brain, *CNC* – commissural neurite bundles of the *VNC, cpnc* – central perikarya of the *VNC, CU* – cuticle, *ECM* – extracellular matrix, *EP* – epidermis, *EXP* – epidermal cell processes, *GA* – giant axons, *GC* – enteral coelom, *LN* – circular neurite bundles, *lpnc* – lateral perikarya of the VNC, *lvtp* – ventrolateral perikarya of the tripartite ventral aggregation, *NE* – neuropile of the lateral brain lobes, *nep* – peripheral perikarya of the lateral brain lobes, *OB* – obturaculum, *OBC* – obturacular coelom, *OP* – opisthosome, *pmp* – posterior median perikarya aggregation, *PNC* – paired strands of the VNC surrounding the ventral ciliary field, *vvtp* - ventral perikarya of the tripartite ventral aggregation, *TR* – trunk, *VNC* – ventral nerve cord, *VT* – vestimentum, *VWF* – collar of the vestimental wings, *VWN* – neurite bundle of the *VW*.

**Fig 2.**
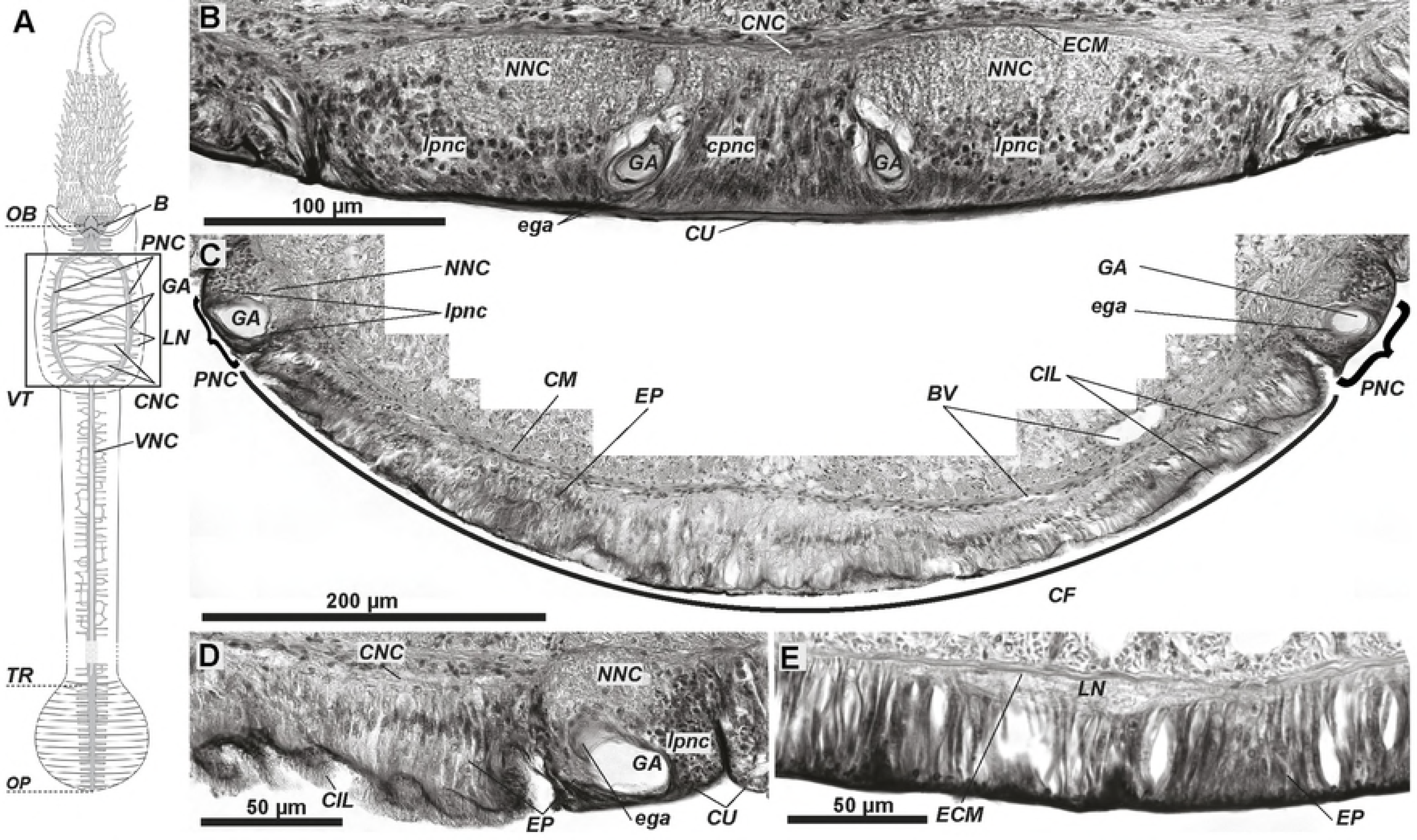
The ventral nerve cord in vestimentum of *Riftia*. A - scheme of the central nervous system which main elements are in grey, and giant axons are in light grey. The frame indicates the area corresponding to histological cross sections (B-E). B - ventral nerve cord just anteriorly to the ventral ciliary field. C - ventral ciliary field (*CF*) surrounded by the paired strands of the ventral nerve cord (*PNC*); line shows the border of the ciliary field, braces show the strands of the ventral nerve cord. D - precise view of the left strand of the ventral nerve cord, commissural neurite bundles connecting the paired strands are seen (*CNC*). E - lateral circular neurite bundles (*LN*) in the epidermis. *B* – brain, *BV* – blood vessels, *CNC* – commissural neurite bundles of the *VNC, CM* – circular musculature, *CF* – ventral ciliary field, *CIL* – cilia, *cpnc* – central perikarya of the *VNC, CU* – cuticle, *ECM* – extracellular matrix, *ega* – cells coating the *GA*, *EP* – epidermis, *GA* – giant axons, *LN* – circular neurite bundles, *lpnc* – lateral perikarya of the *VNC, NNC* – neuropile of the VNC, *OB* – obturaculum, *OP* – opisthosome, *PNC* – paired strands of the VNC surrounding the CF, *TR* – trunk, *VNC* – ventral nerve cord, *VT* – vestimentum.

**Fig 3.**
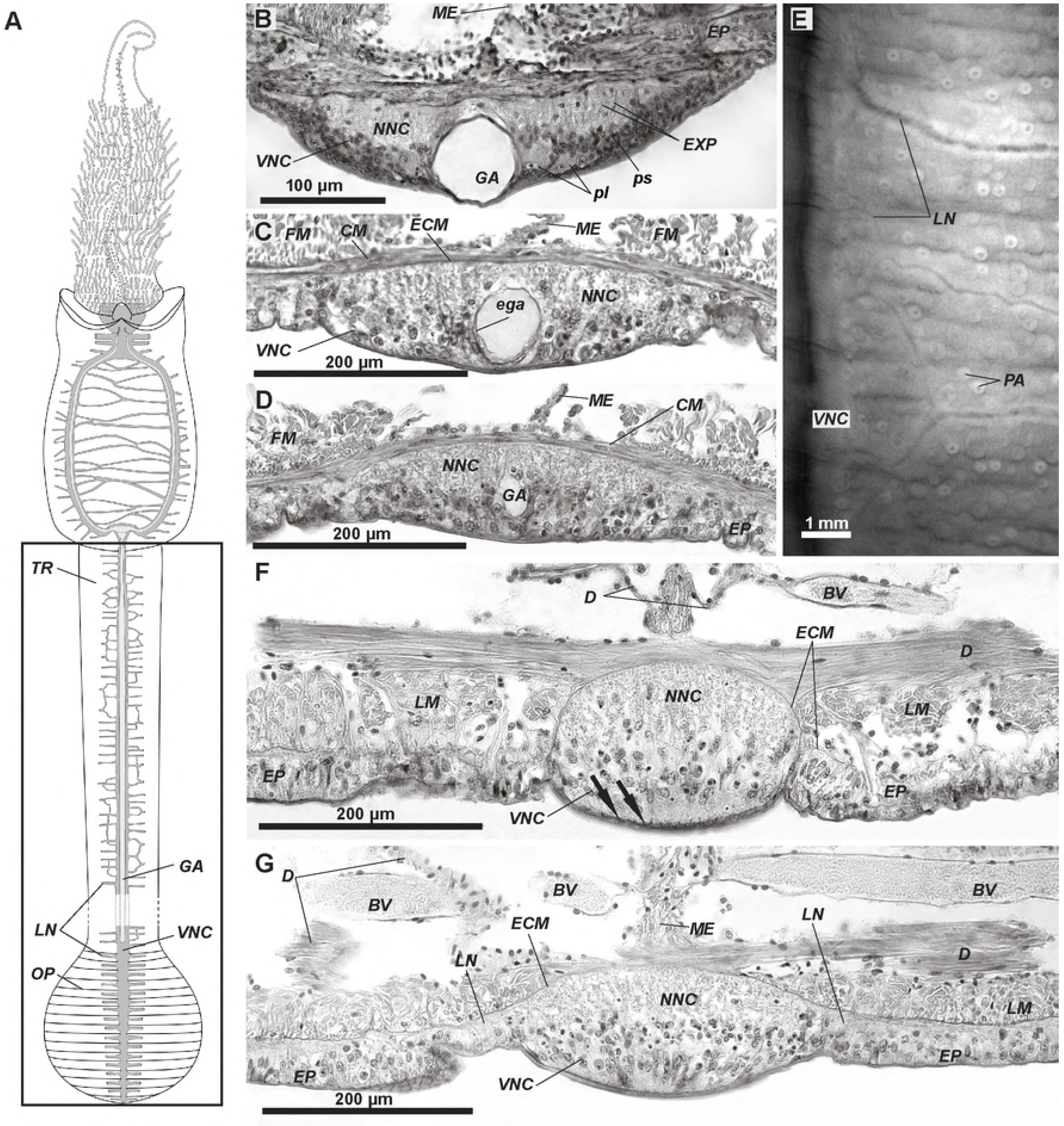
The ventral nerve cord in trunk and opisthosome of *Riftia*. A - scheme of the central nervous system which main elements are in grey, and giant axons are in light grey. The frame indicates the area corresponding to histological sections (B-D, F, G) and light miscroscopical image (E). B – the ventral nerve cord (VNC) structure in the anterior trunk. C, D - VNC structure in the midtrunk and posterior trunk, respectively; note the reduction of the giant axon diameter. E - lateral neurite bundles branching and making anastomoses in the trunk epidermis. F-E - VNC in the middle and posterior part of the opisthosome. Arrows in (F) show the cuticular folds between cell borders. *ECM* – extracellular matrix, *BV* – blood vessels, *CM* – circular muscles, *D* – dissepiments, *ega* – cells coating the *GA*, *EP* – epidermis, *EXP* – epidermal cell processes, *GA* – giant axons, *FM* – featherlike longitudinal muscles, *LG* – longitudinal lateral grooves, *LM* – longitudinal muscles, *LN* – circular neurite bundles, *lpnc* – lateral perikarya of the *VNC, ME* – mesenterium, *NNC* – neuropile of the VNC, *OP* – opisthosome, *PA* – cuticular plaque papillae, *pl* – large perikarya, *ps* – small perikarya, *TR* – trunk, *VNC* – ventral nerve cord.

**Fig 4.**
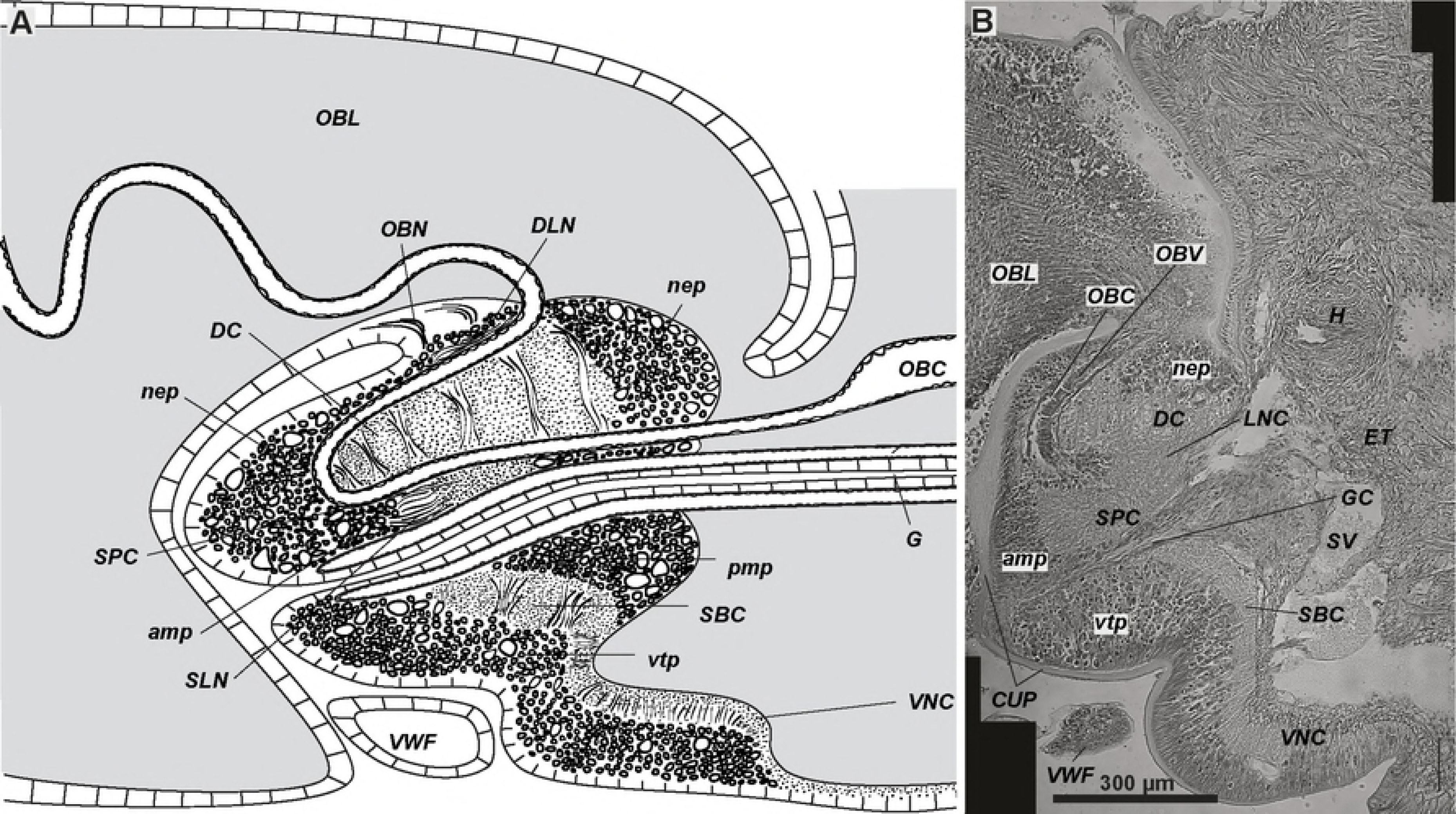
Brain of juvenile *Riftia* with a gut rudiment. A - scheme of the sagittal section of the vestimentiferan brain which consists of supraesophageal and subesophageal ganglia. B - parasagittal section of the 8 mm long juvenile, the gut rudiment passes through the vestimentiferan brain. *amp* – anterior median aggregation of perikarya, *CUP* – cuticle schild, *DC* – dorsal commissure, *DLN* – dorsal area of the longitudinal bundles, *ET* – excretory tree, *G* – gut lumen, *GC* – enteral coelom, *H* – heart, *LNC* – lateral connectives, *nep* – peripheral perikarya of the lateral brain lobes, *OBC* – obturacular coelom, *OBL* – obturacular lobes, *OBN* – obturacular neurite bundles, *OBV* – obturacular blood vessels, *pmp* – posterior median perikarya aggregation, *SBC* – subenteral commissure, *SLN* – supraenteral longitudinal neurite bundles, *SPC* – supraenteral commissure, *SV* – *sinus valvatus, vtp* – tripartite ventral aggregation of perikarya, *VNC* – ventral nerve cord, *VWF* – collar of the vestimental wings.

Ventral brain lies in the anteriormost vestimentum (Figs 1A, 4A-B). There are two brain lobes forming heart-like shape on transverse sections (Figs 5–7, S1-S3 Figs). Dorsal furrow between the brain lobes encloses the **obturacules’** bases (*OBL*, Figs 4A, 5-6, S1 Fig). Posteriorly the **excretory tree** is adjacent to the brain (*ET* Fig 4B). The whole brain lies inside the **epithelium**, and there are no basal laminae separating brain from epidermis (*EP*, Figs 4–7, S1-4). **Cuticle schild** protects the apical surface of the brain (*CUP*, Figs 4B, S1-3 Figs). **Collar of vestimental wings** shelters the ventral brain from outside (*VWF*, Fig 1B, 4). The 80 mm long specimen has a brain of 1 mm in height and length, 2 mm in width. **Undifferential tentacle lamellae** lie on the dorsal and lateral surface of the brain (*LR*, Figs 5-7, S1-S3A, S5 Figs). Posterior brain has mainly dorsal lamellae (Figs 6–7, S3A Fig), whereas in anterior brain the tentacle lamellae occupy dorsal surface as well as descend to the lateral and ventrolateral surfaces (Figs 5, S1, S2 Figs).

**Fig 5.**
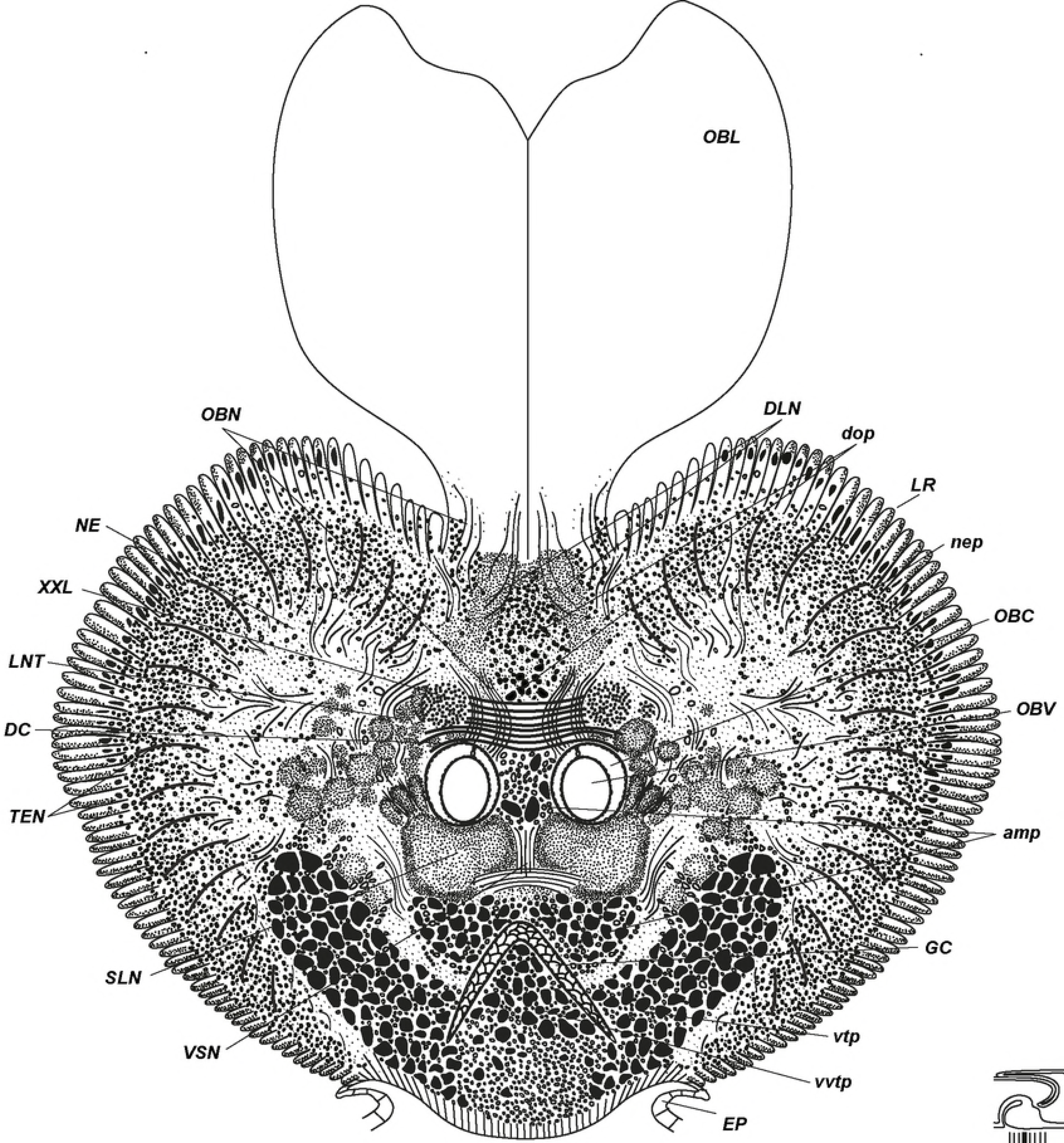
Anterior brain organization of *Riftia*. Scheme of histological cross section based on anterior brain sections of 79 mm long specimen (see S1 Fig). Level of the section shown at the diagram at the right lower coner. *amp* – anterior median aggregation of perikarya, *DC* – dorsal commissure, *DLN* – dorsal area of the longitudinal bundles, *dop* – dorsal aggregation of perikarya, *EP* – epidermis, *GC* – enteral coelom, *LNT* – longitudinal nerve tracts projecting from the *VNC* into the brain, *LR* – undifferential tentacle lamellae, *NE* – neuropile of the lateral brain lobes, *nep* – peripheral perikarya of the lateral brain lobes, *OBC* – obturacular coelom, *OBL* – obturacular lobes, *OBN* – obturacular neurite bundles, *OBV* – obturacular blood vessels, *SLN* – supraenteral longitudinal neurite bundles, *TEN* – neurite bundles of tentacles (palps), *VSN* – vertical supraenteral neurite bundles, *vtp* - tripartite ventral aggregation of perikarya, *vvtp* - ventral perikarya of the *vtp, XXL* – pair of prominent bundles of large longitudinal nerve tracts (part of *LNT*).

**Fig 6.**
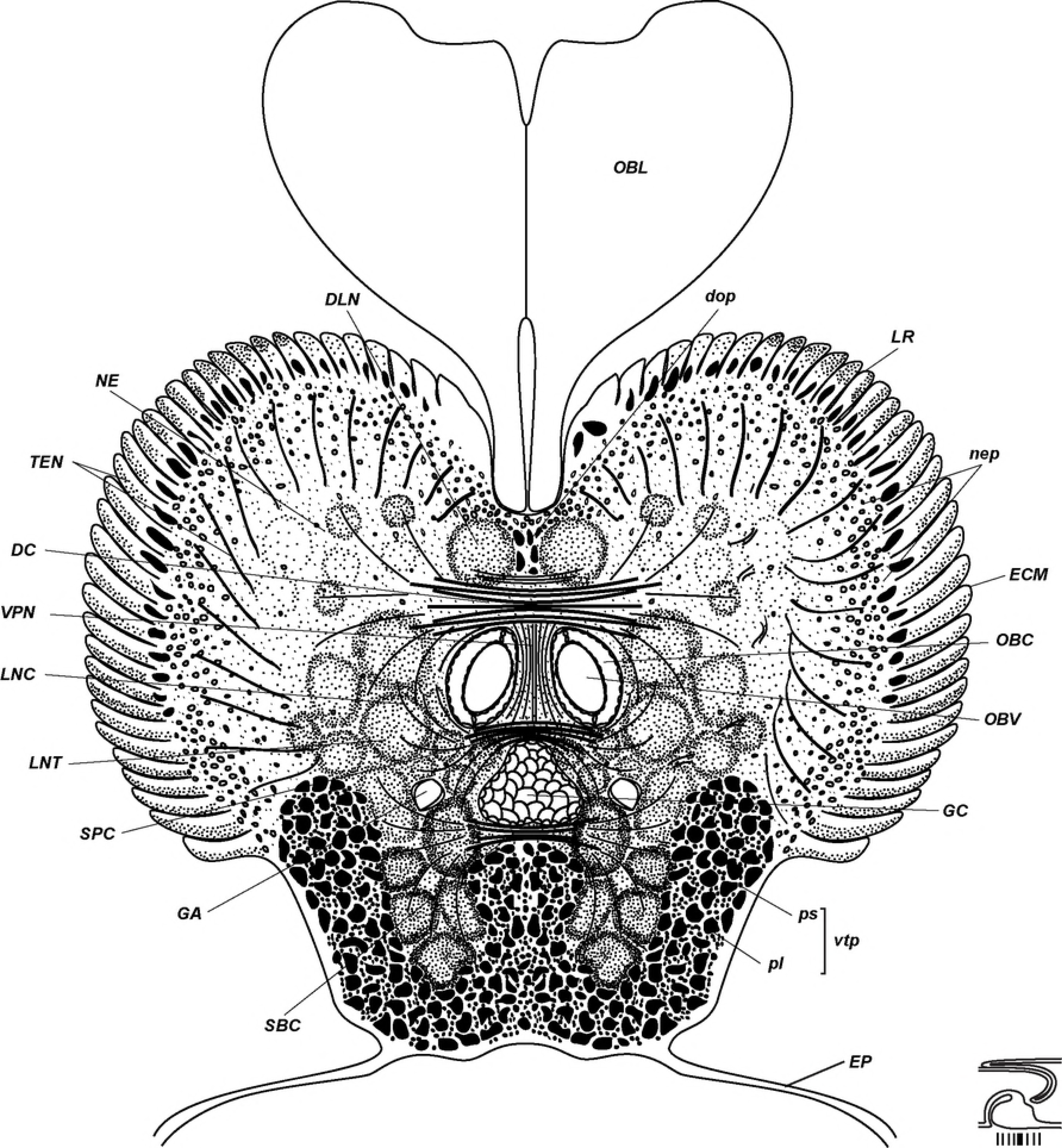
Middle brain organization of *Riftia*. Scheme of cross section based on midbrain histological sections of 79 mm long specimen (see S2 Figure). Level of the section shown at the diagram at the right lower coner. *DC* – dorsal commissure, *DLN* – dorsal area of the longitudinal bundles, *dop* – dorsal aggregation of perikarya, *EP* – epidermis, *GA* – giant axons, *GC* – enteral coelom, *ECM* – extracellular matrix, *LNC* – lateral connectives, *LNT* – longitudinal nerve tracts projecting from the *VNC* into the brain, *LR* – undifferential tentacle lamellae, *lvtp* – ventrolateral perikarya of the *vtp, NE* – neuropile of the lateral brain lobes, *nep* – peripheral perikarya of the lateral brain lobes, *OBC* – obturacular coelom, *OBL* – obturacular lobes, *OBV* – obturacular blood vessels, *pl* – large perikarya, *ps* – small perikarya, *SBC* – subenteral commissure, *SPC* – supraenteral commissure, *TEN* – neurite bundles of tentacles (palps), *vtp* - tripartite ventral aggregation of perikarya, *VPN* – posterior vertical median bundles.

**Fig 7.**
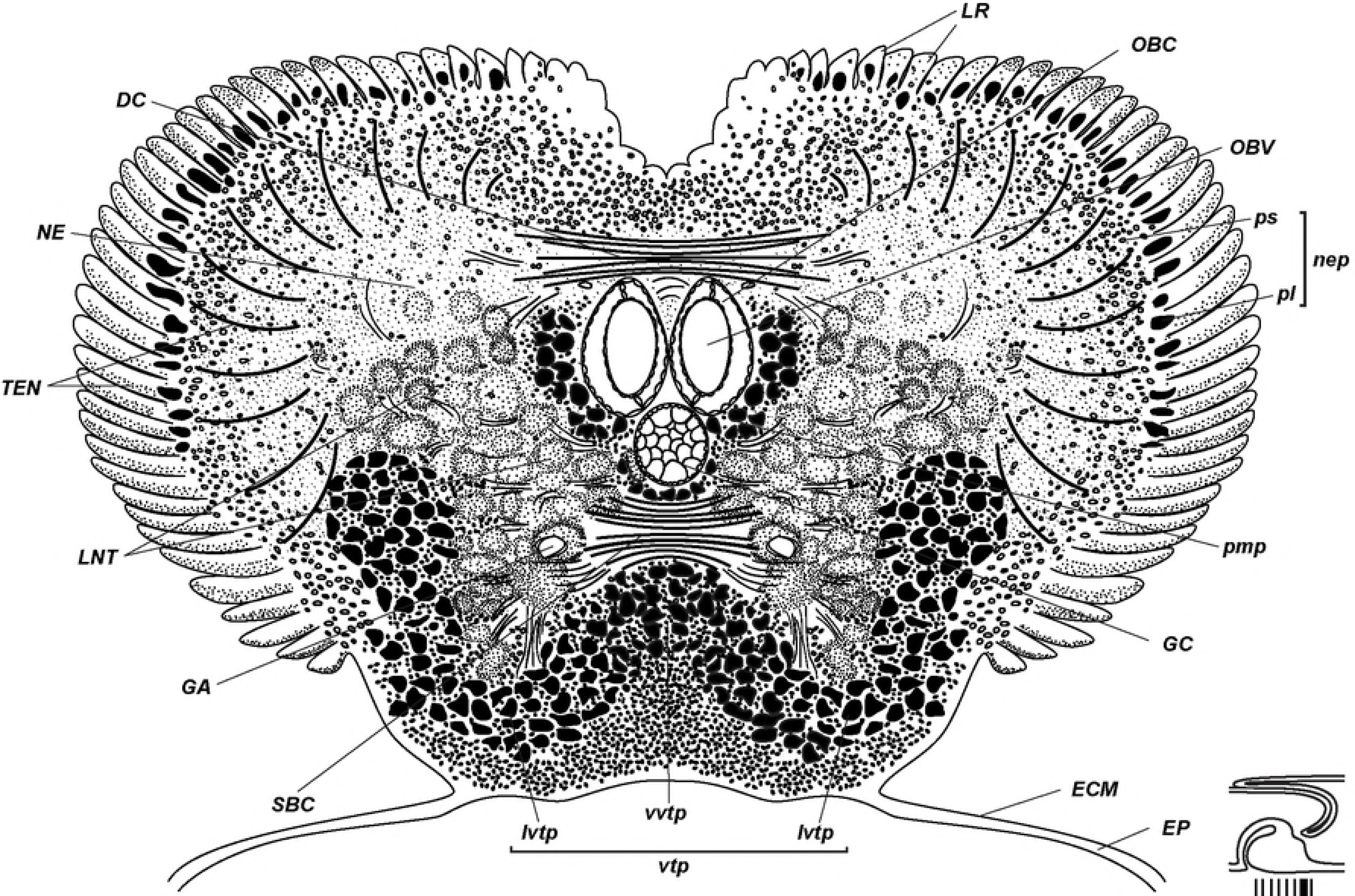
Posterior brain organization of *Riftia*. Scheme of cross section based on posterior brain histological sections of 79 mm long specimen (see S3 Figure). Level of the section shown at the diagram at the right lower coner. *DC* – dorsal commissure, *ECM* – extracellular matrix, *EP* – epidermis, *GA* – giant axons, *GC* – enteral coelom, *LNT* – longitudinal nerve tracts projecting from the ventral nerve cord into the brain, *LR* – undifferential tentacle lamellae, *lvtp* – ventrolateral perikarya of the *vtp, NE* – neuropile of the lateral brain lobes, *nep* – peripheral perikarya of the lateral brain lobes, *OBC* – obturacular coelom, *OBV* – obturacular blood vessels, *pl* – large perikarya, *pmp* – posterior median perikarya aggregation, *ps* – small perikarya, *SBC* – subenteral commissure, *TEN* – neurite bundles of tentacles (palps), *vtp* - tripartite ventral aggregation of perikarya, *vvtp* - ventral perikarya of the *vtp*.

Three coelomic channels pass through the brain tissue: pair of **obturacular coeloms** with blood vessels and unpaired **enteral coelom** (*OBC, GC*, Figs 4–7, S6 Fig). In juvenile undivuduals the enteral coelom comprises **gut rudiment** (G, Fig 4A). In larger specimens the enteral coelom is occupied with mesenchymal cells (Figs 6). In anterior brain the enteral coelom has «Λ» shape of transverse profile (Figs 5). Oral siphon is preserved in juvenile *Riftia* having 34 mm in length, and the intestine rudiment remains in the coelomic channel running through the brain, in individuals having 79 mm in length. In larger individuals the only coelomic channel remains.

Ventral nerve cord (*VNC*) connects to the brain *via* longitudinal neurite bundles (Figs 1B-E). Anteriorly to the **ventral ciliary field** (*CF*) the VNC splits into a **pair of strands** (*PNC*) connected to each other with transverse neurite bundles (Fig 2A-D). The strands surround the ventral ciliary field (Figs 2B-C). The strands fuse into a single VNC at the border of the vestimentum and trunk and extend along ventral midline till the end of the body (Fig 3A). The width of the prominent VNC can reach up to 1 mm in a specimen of 808 mm long (Fig 3B). Its width decreases to posterior trunk (Figs 3C-D).

The VNC is lying inside the epidermis (Fig. 1E). The epidermal cells have a wide apical part adjacent to cuticle and basal process to the layer of the ECM (Fig. 1C).

Apically thick cuticular layer (CU) protects VNC, especially in the anteriormost part (Fig 1C). In opisthosome the cuticle protecting VNC makes folds between the apical parts of epidermal cells (arrows, Fig 3F).

### Dorsal brain structures

Brain of *R. pachyptila* consists of dorsal and ventral parts divided by position of the enteral coelom (Figs 4-8).

**Fig 8.**
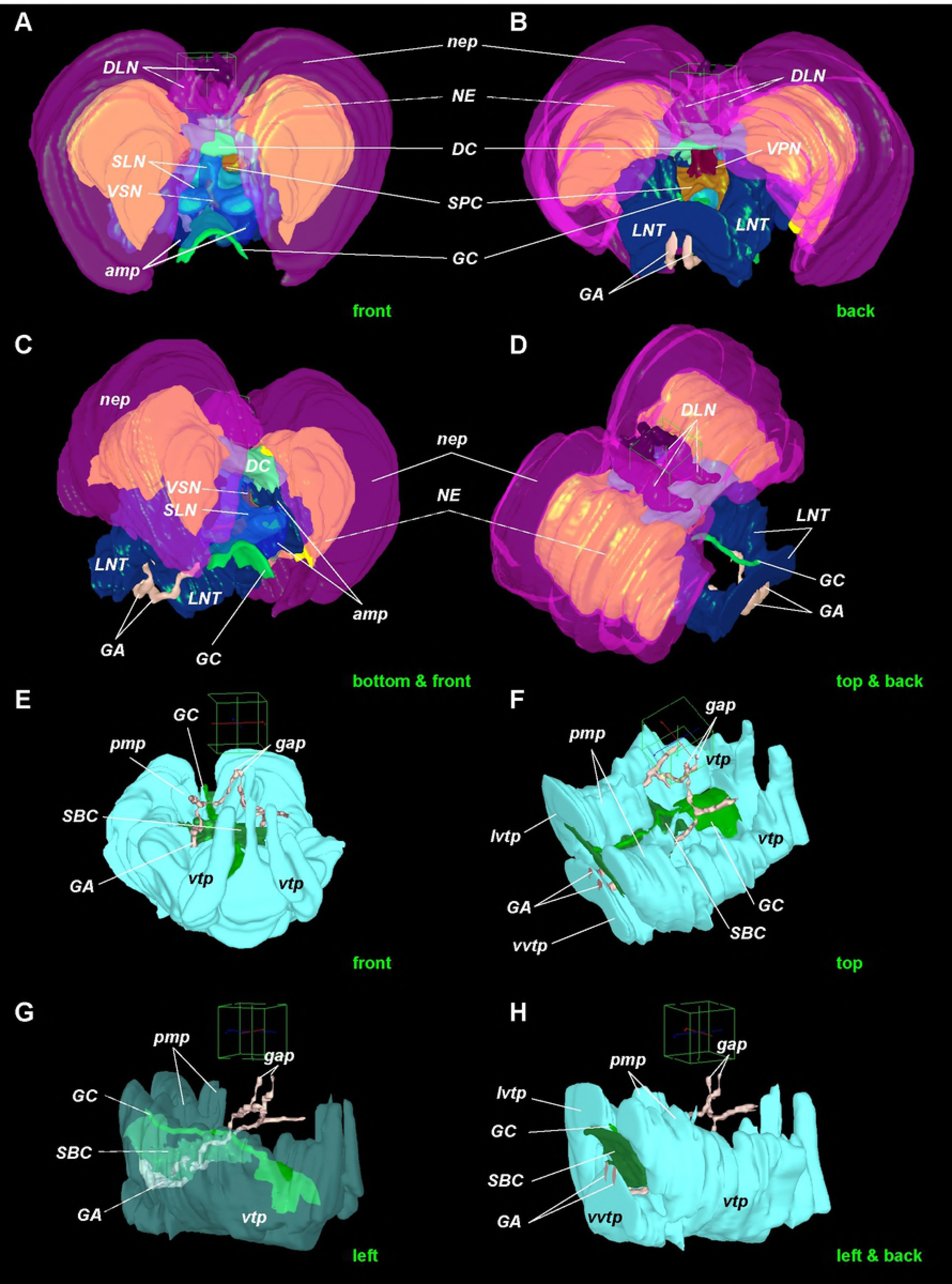
Supra-and subesophageal ganglia in *Riftia*. 3D models of *Riftia* brain. A-D - supraesophageal neuronal elements. E-H - subesophageal neuronal elements in *Riftia* brain. View sides shown at the right lower corners of each images. Cube side is 255 μm. Dashed lines point neural elements under transparent structures. *amp* – anterior median aggregation of perikarya, *DC* – dorsal commissure, *DLN* – dorsal area of the longitudinal bundles, *GA* – giant axons, *gap* – giant perikarya, *GC* – enteral coelom, *LNT* – longitudinal nerve tracts projecting from the ventral nerve cord into the brain, *lvtp* – ventrolateral perikarya of the *vtp, NE* – neuropile of the lateral brain lobes, *nep* – peripheral perikarya of the lateral brain lobes, *pmp* – posterior median perikarya aggregation, *SBC* – subenteral commissure, *SPC* – supraenteral commissure, *SLN* – supraenteral longitudinal neurite bundles, *VPN* – posterior vertical median bundles, *VSN* – vertical supraenteral neurite bundles, *vtp* - tripartite ventral aggregation of perikarya, *vvtp* - ventral perikarya of the *vtp*.

Most part of the dorsal brain occupied by paired areas of **neuropile of the lateral brain lobes** (*NE*) in the thickness of which there are many perikarya (Figs 5–8, S1-S3, S7A-C’, S8 Figs). Numerous radial tentacle neurite bundles extend from the neuropile of the lateral brain lobes to the bases of the tentacle lamellae, these are **tentacle neurite bundles** (*TEN*, Figs 5–6, S1-S3, S5 Figs). Each lamella represents the thin fold of the epidermis (Figs S5 A-D). Lamellae are closely adjacent to each other, and **epidermis of external lamellae wall** (*OEP*) is flattened, **epidermis of the internal wall** (*IEP*) is thicker and contain the basiepithelial tentacle neurite bundles (S5A Fig).

Neuropiles of right and left brain lobes are connected by thick extended **dorsal commissure** (*DC*, Figs 4, 5-7, 8A-C, 9, S1-S3, S7A-H Figs). It lies over the paired obturacular coelomic channels and adjacent to their loops anteriorly (S8F-G Figs). Transverse neurite bundles included in the dorsal commissure are divided into two almost equal parts: anterior and posterior commissures (Figs 9A, B, E, S8F, G Figs). Both dorsal commissures (anterior and posterior) of large specimen are structured in dorso-ventral direction and comprises of several layers of neurite bundles which are visible at the transverse section (up to 5 levels in 79 mm long specimen, S2A Fig). Up to 9-11 ventro-dorsal vertical bundles go through the dorsal commissure clearly visible at sagittal and parasagittal sections (Fig 4A).

**Fig 9.**
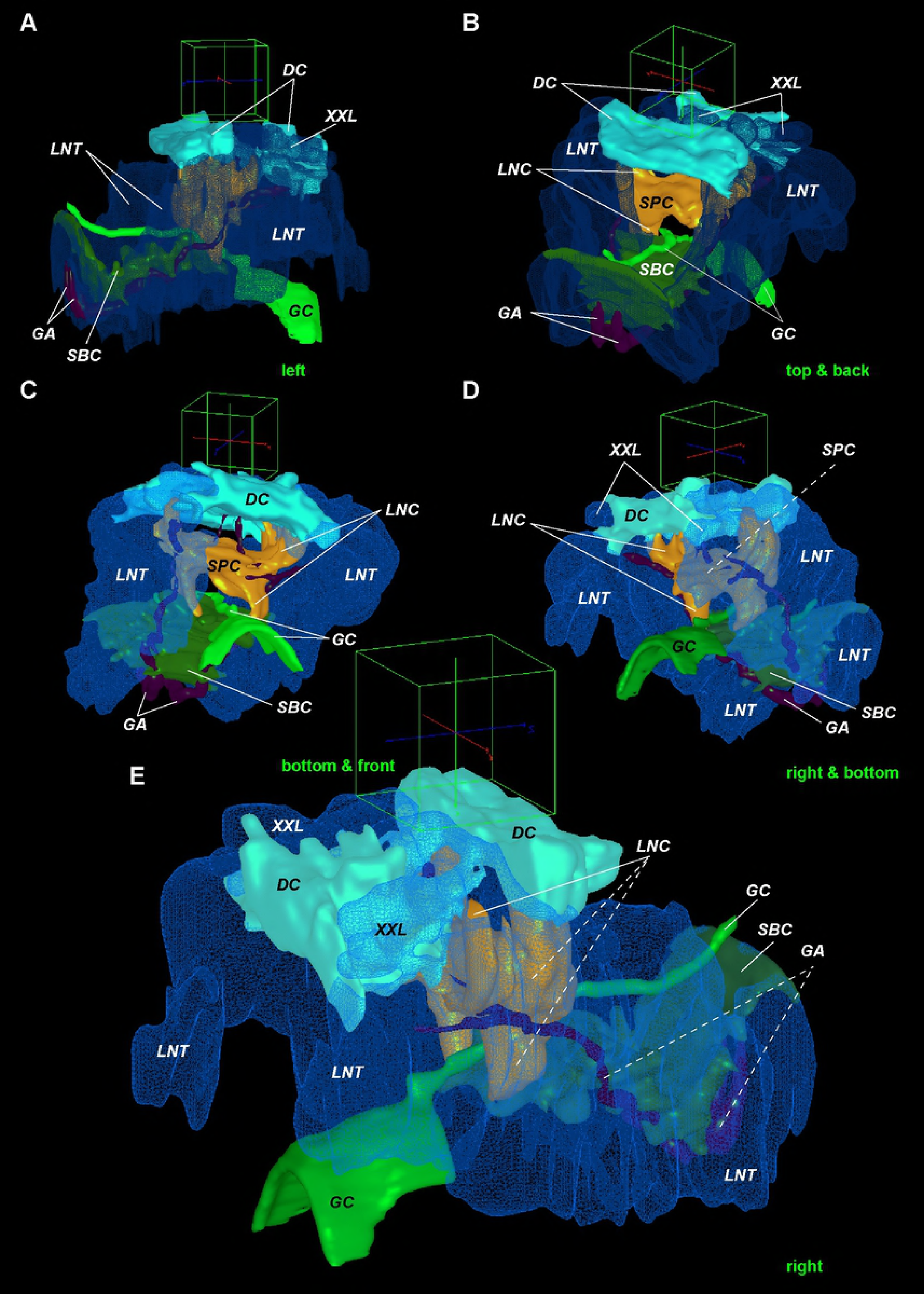
Longitudinal nerve tracts and main commissures in *Riftia* brain. 3D models of *Riftia* brain. A-E - main commissures (dorsal, *DC*, supra-, *SPC*, and subesophageal, *SBC*) and longitudinal nerve tracts (*LNT*). The latter is homologous to circumesophageal connectives in another annelids’ brain. View sides shown at the right lower corners of each images. Cube side is 255 μm. Dashed lines point neural elements under transparent structures. *DC* – dorsal commissure, *GA* – giant axons, *GC* – enteral coelom, *LNC* – lateral connectives, *LNT* – longitudinal nerve tracts projecting from the ventral nerve cord into the brain, *SBC* – subenteral commissure, *SPC* – supraenteral commissure, *XXL* – pair of prominent bundles of large longitudinal nerve tracts (part of *LNT*).

Two pairs of **obturacular neurite bundles** (*OBN*) extend from the dorsalmost area of the brain from the anterior dorsal commissure to the bases of obturacular lobes (Figs 5, 10A, S1, S9A-F). Each pair of obruracular bundles (left and right) gives rise neurite bundles in the epidermis of inner and outer sides of the obturacular lobes. In that area neurite bundles run vertically, then in the dorsal part of obturacules move in posterior-anterior direction.

In the midbrain there is a weak **supraenteral commissure** (*SPC*) running over the enteral coelomic channel, but under the obturacular channels (Figs 4A, 9B-D, 10B, C, S3A, S8D-E, S9J, S10B, D Figs). In anterior brain two prominent **supraenteral longitudinal neurite bundles** (*SLN*) directs backward from the anteriormost brain and disintegrate into separate small bundles at the level of the supraenteral commissure (compare Figs 5&6, Figs 8A, S1, S2, S8, S10A, C, D Figs). Supraenteral longitudinal neurite bundles are connected to each other *via* **vertical supraenteral neurite bundles** (*VSN*) which have inverted «Ύ»-like shape (Figs 5, S2B, S8A-C Figs). Moreover, the vertical bundles join different parts of fibers of an **anterior median aggregation of perikarya** (*amp*): vertical fibers connect dorsal and ventral parts of the aggregation, as well as transverse fibers connect left and right halves of the aggregation (Figs 4, 5, S2, S8 Figs).

**Fig 10.**
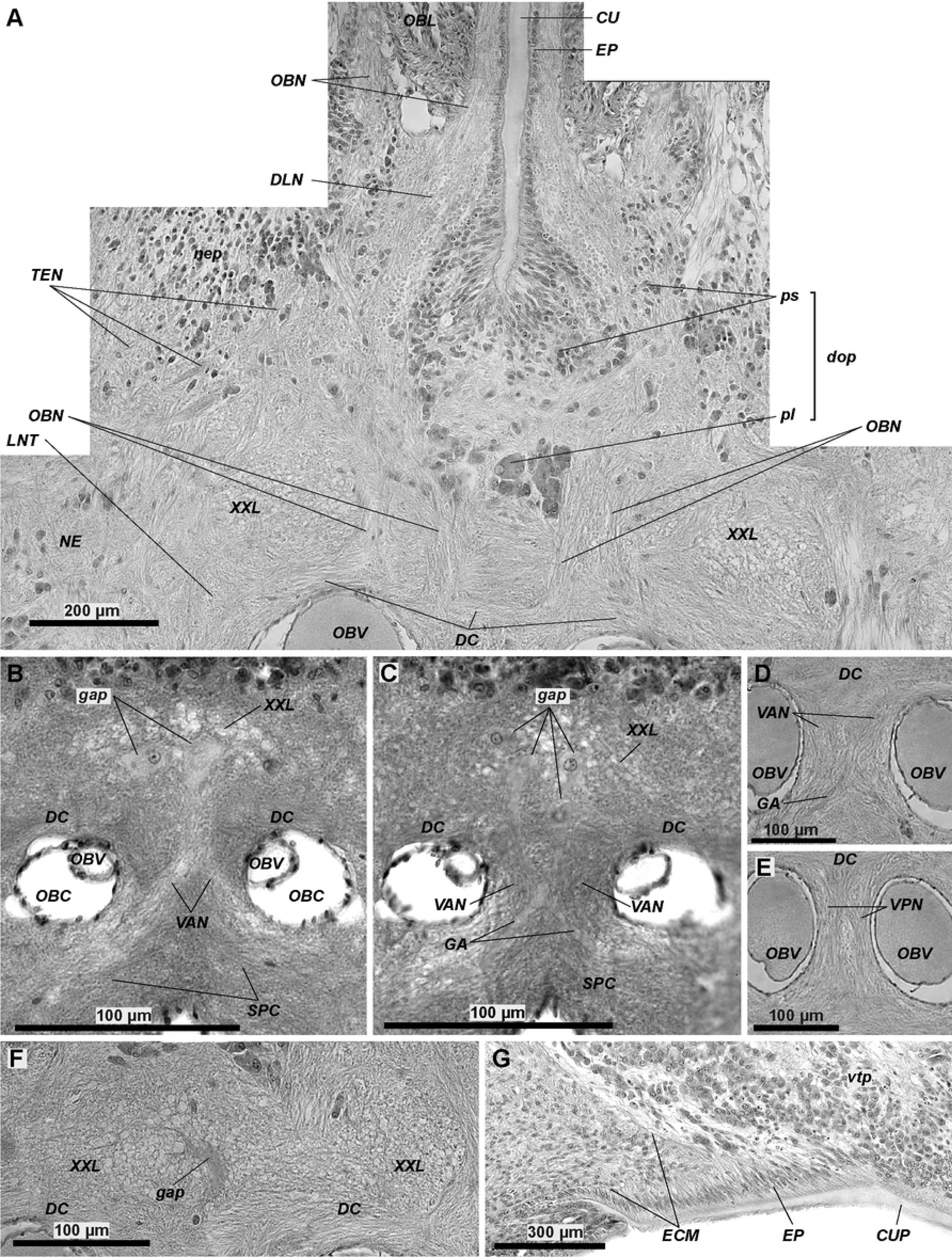
Histological details in the anterior brain of *Riftia*. A - obturacular neurite bundles (*OBN*) connecting with the dorsal commissure (*DC*). B, C - giant perikarya with clear nuclei in juvenile brain. D - anterior vertical median bundles (*VAN*) comprising of giant axons (*GA*). E - posterior vertical median bundles (*VPN*) with no giant axon. F - giant perikarion degrading in brain of 79 mm long male. *G* - cuticular plate protecting the brain (*CUP). ECM* – extracellular matrix, *CU* – cuticle, *CUP* – cuticle schild, *DC* – dorsal commissure, *DLN* – dorsal area of the longitudinal bundles, *dop* – dorsal aggregation of perikarya, *EP* – epidermis, *GA* – giant axons, *gap* – giant perikarya, *LNT* – longitudinal nerve tracts projecting from the *VNC* into the brain, *NE* – neuropile of the lateral brain lobes, *nep* – peripheral perikarya of the lateral brain lobes, *OBC* – obturacular coelom, *OBL* – obturacular lobes, *OBN* – obturacular neurite bundles, *OBV* – obturacular blood vessels, *pl* – large perikarya, *ps* – small perikarya, *SPC* – supraenteral commissure, *TEN* – neurite bundles of tentacles (palps), *vtp* - tripartite ventral aggregation of perikarya, *VPN* – posterior vertical median bundles, *VAN* – anterior vertical median bundles, *XXL* – pair of prominent bundles of large longitudinal nerve tracts (part of *LNT*).

On the dorsalmost side of the midbrain (close to ECM layer) there is a pair of **dorsal areas of the longitudinal bundles** (*DLN*, Figs 4A, 5, 6, 8A-D, 10A, S1, S2, S3A, S9A-C). They start from a **dorsal aggregation of perikarya** (*dop*) in the midrain (Figs 5, 10A, S1, S2 Figs) and lie along the dorsal groove of the brain. The dorsal areas of the longitudinal bundles expand widely along the site of the groove untill the place of obturacules enter the brain.

Short **anterior vertical median bundles** (*VAN*) pass between the obturacular coeloms in the midbrain (Figs 10B-D, S10 Fig). They extend ventro-dorsally between the supraenteral commissure and the the roots of anterior dorsal commissure. Anterior vertical median bundles comprise of the crossing neurite bundles: the neurite bundles originating from the right side of the supraenteral commissure extend to the left side of the dorsal commissure, and vice versa.

Posteriorly to the anterior vertical median bundles there are **posterior vertical median bundles** (*VPN*, Figs 6, 8B, 10E, S3A, S10 Figs). They do not contain any crossing bundles and connect the supraenteral commissure and the posterior dorsal commissure.

**Periferic perikarya of the lateral brain lobes** (*nep*, Figs 4-7, 8A-D, S1-S3, S5A, S6A, C, E, S7I, J, S9A Figs) are represented by two layers: inner layer of small perikarya, 5 μm (*ps*), and outer layers of big ones (*pl*), 20 μm (Fig 7, S3A, S5A Figs). In juvenile specimens having lower number of tentacle lamellae the small perikarya are grouped into distinct lobules which correspond to the tentacle lamellae. In bigger specimens having higher number of tentacle lamellae arrangement of small perikarya are even. In anterior part of the brain the periferic zone of perikarya expands significantly and covers laterally a **tripartite ventral aggregation of perikarya** (*vtp*, Figs 5–7, S1-S3, S7I, J, Figs, more about *vtp* read below).

In the dorsal groove of the anterior brain there is a **dorsal aggregation of perikarya** (*dop*) which lies in the inner sides of obturacules entering the brain (Figs 5, 6, 10A, S1, S2, S3A Figs). It contains two layers of perikarya: in contrast to perifiric perikarya there are inner big perikarya and outer small ones (Figs 5, 10A).

The **anterior median aggregation of perikarya** (*amp*) is the most anterior symmetrical accumulation of big somata (Fig 4, 5, 8A, C, S1, S2, S8 Figs). It is adjacent dosally to the enteral coelomic channel.

### Ventral brain structures

In the ventral brain, under the enteral coelomic channel, there is a main **subenteral commissure** (*SBC*, Fig 4, 6, 7, 8E-H, 9, S3, S8D-G Figs) which is a continuation of the transverse neurites in the ventral nerve cord (*CNC*, Fig. 2B).

The most neuropile of the ventral brain is occupied by paired prominent **longitudinal nerve tracts** (*LNT*, Figs 1B-D, 6, 7, 8B-D, 9, 10A, S1-S3, S7, S9B-J Figs) which are continuations of nerve fibers from the ventral nerve cord (Figs 1B-D). As the longitudinal nerve tracts come into the brain, each of them lies around three coelomic channels and gradually rises to the dorsal side of the brain (Figs 5-7).

In the dorsal brain the tracts contain a **pair of large bundles** of thick fibers, 6-11 μm in diameter of a fiber (*XXL*, Figs 5, 9A, B, E, 10A-C, F, S2B, S7D, E, G, S9B, C Figs). Anteriorly the prominent bundles fall apart into several smaller bundles which disintegrate in the neuropile of the lateral brain lobes (S7A’, B’, D, G Figs).

Neurites of the neuropile of the lateral brain lobes (*NE*) originate from the longitudinal nerve tracts (Figs 5-7, 8B-D, S1-S3, S7 Figs).

In the anterior dorsal brain, the longitudinal nerve tracts are connected to each other *via* the dorsal commissures (*DC*) over the obturacular coeloms (Figs 5, 9, S1, S2, S7A’-H, S9B-C Figs) and *via* the supraenteral commissure (*SPC*) under the obturacular coeloms (Figs 6, 9B-D, S3A Fig). In the ventral brain the pair of the longitudinal nerve tracts are binded by subenteral commissure (*SBC*, Figs 6, 7, 9, S7B’, C’, G Figs).

The ventralmost part of the brain, under the enteral coelom, is occupied with the **tripartite ventral aggregation of perikarya** (*vtp*, Figs 4–7, 8E-H, 10G, S1, S2A, S6A-D, S7A-C, I, J, S8D, E Figs) comprising of small and big perikarya (Fig 6). On transverse sections it is divided in three lobes: ventral and two ventrolateral ones (*vvtp, lvtp*, Figs 7, S2B, S3, S6 Figs). In the posterior brain lobes enter the brain neuropile significantly (Figs 6, 7, S3). In anterior brain the unpaired ventral lobe adjoins the ventral side of the enteral coelomic channel (Figs 5, 8F, G, S1, S2, S6D-F Figs). In the posterior brain two groups of big perikarya, **posterior median perikarya aggregations** (*pmp*), extend from the tripartite aggregation forward and lie along the left and right sides of three coelomic channels (Figs 7, 8E-H, S3B, S6F Figs).

### Giant perikaria and axons

**Giant axons** run in the middle and posterior brain parts (*GA*, Figs 6, 7, 8B-H, 9, S3, S9G-L, S10B-G Figs). In juvenile and male specimens, we found two pairs of dorsal giant perikarya (*gap*) lying in the dorsal neuropile of the dorsal commissure and the longitudinal nerve tracts (Figs 10B, C, F). Nuclei as well as nucleoli remain in the **giant perikarya** of juvenile, but not in male specimen (Figs 10B, C). Axons of the dorsal giant perikarya run ventrally as part of the crossing **anterior vertical median bundles** (*VAN*, Figs 10B-D, S10 Fig).

3D-modelling of studied juveniles revealed two pairs of lateral branches of giant axons in lateral neuropiles of longitudinal nerve tracts which do not have giant perikarya (S9G, L, S10E-G Figs). Perhaps in younger specimens they remain. In the posterior brain the giant axons extend inside the longitudinal nerve tracts and continue inside the neuropile of the ventral nerve cord (Figs 9, S9G-J Figs). Transversally the giant axon represents the 20-25 μm round profile with light cytoplasm and enveloped by flattened cells with dark nuclei (S3 Fig).

### Ventral nerve cord

In vestimentum neuropile of the paired ventral nerve cord (VNC) consists of two lateral longitudinal nerves (*LNT*, Figs 1B-D) connected *via* transverse (commissural) neurite bundles (*CNC*, Figs 1B, D, 2B). Pair of giant axons lies in the central part of VNC (Figs 1A-D). Numerous small perikarya form two lateral and one central accumulations (*lpnc*, cpnc, Figs 1C, D) which are continuations of the ventral tripartite aggregation of the ventral brain (*vvtp*, Figs 1B, C).

Around the ventral ciliary field each strand of *PNC* contains the epidermal cells, basal neuropile, apical perikarya, and single fiber of the giant axon envelopped with the coating cells (*ega*) (Fig 2D). Most perikarya lie externally to the giant axon in each strand. The ciliary field consists of columnar ciliary epidermal cells (Fig 2C). In their basal parts there are commissural neurite bundles (*CNC*) which make a net and connect the strands with each other (Figs 2A, B, D).

In trunk the VNC has permanent diameter, neuropile has no swellings and separated by the giant axon into two longitudunal strands (Figs 3A-D). The epidermal cells’ processes extend to the ECM inside the neuropile (*EXP*, Fig 3B). The VNC perikarya spread along left and right sides of the giant axon (Figs. 3B-D). There are small (3,5 μm) and big (20 μm) perikarya (*ps, pl*, Fig 3B). Giant axon extends to the border of the trunk and opisthosome (Fig 3D).

In opisthosome an arragement of the apical somata and basal neuropile of the VNC is the same as in the rest body (Figs 3F, G). There is no giant axon, all perikarya are small.

### Segmental nerve bundles

In the anteriormost vestimentum several thick transverse **lateral neurite bundles** part off the ventral nerve cord (*LN*, Fig 1A). We found 3 pairs of them in 16 mm long specimen. The first pair, the most prominent one, directing to the anterior collar, is **neurites of vestimental wings** (*VWN*, Figs 1A, B). At the level of the ciliary field, many irregular bundles part off the lateral neuropile of the VNC strands and extend into the epidermis of the vestimental wings (Figs 2A, E). Transverse neurite bundles come off the single VNC in the trunk. They intensively branch and make anastomoses (Fig 3E). In a 79 mm long specimen, lateral bundles part each 100 μm off the cord, thus there are 350-360 pairs of bundles in a trunk. In each opisthosomal segment a pair of lateral bundles leaves the neuropile of the VNC (Fig 3A, compare F&G).

## Discussion

### Ventral nerve cord in Vestimentifera

To date described species of vestimentiferans have uniform structure of the ventral nerve cord, except the length of giant axons and organization of perikarya aggregations in trunk [7,10–12,15,53,54]. The ventral nerve cord in *Ridgeia piscesae* and *Lamellibrachia satsuma* comprises of central neuropile and two lateral strands of perikarya, thus showing somewhat paired structure [10,12], whereas in *Riftia* (present study) and *Oasisia alvinae* there are single layers of apical perikarya and basal neuropile [11]. Also, in *O. alvinae* median groove was found to run along the midline of the ventral nerve cord in opisthosome [11].

Pair of giant axons extended from the pair of giant perikarya was found in vestimentiferans *Ridgeia, Riftia, Oasisia, Lamellibrachia* [5,10,11,13]. Giant axons terminate at different levels in trunk nerve cord: in *L. luymesi*, giant axons terminate in the anterior part of the trunk segment [53], in *L. barhami* extend a little further back [55], in *R. piscesae, O. alvinae* and *Riftia* they extend up to the border between trunk and the first opistosome segment [10,11]. Earlier a pair of giant perikarya was found to be retained in juveniles of *R. piscesae* and *O. alvinae* in the mid-dorsal part of the brain [10,11,13,15]. We found two pairs of giant neurons in juveniles in the dorsal commissure of *Riftia* (Fig 10 B, C). Besides, the lateral branches of giant axons (S10E-G Figs) indicate the possible presence in earlier stages two pairs of giant perikarya in the lateral areas of the neuropile. Thus, each giant fiber in *Riftia* is a product of the fusion of at least four pairs of axons.

### Ventral nerve cord in Siboglinidae

Siboglinids have intraepidermal ventral nerve cord along which most of perikarya evenly dispersed [9,11,12,19,20,22,23,26–28]. All siboglinids have paired structure of the ventral nerve cord. First, the ventral nerve cord of vestimentiefrans and frenulates have paired structure in vestimentum and forepart, respectively. Second, there is a pair of giant axons in vestimentiferans and large frenulates. Third, in frenulates, vestimentiferans and *Sclerolinum* the ventral cord bifurcates into two strands around the ventral ciliated field. In female *O. priapus*, the only *Osedax* species with the ventral ciliary field, pair of the ventral cords adjoins the ciliary field. Fourth, in *Osedax* species (females and males) there is an obvious pair of widely separated strands of the ventral nerve cord in trunk [9,19,20,22,26–28].

The ventral ciliary field which is unique stucture conserved in all adult siboglinids lies in the anterior worm part: in trunk of frenulates, in vestimentum of vestimentiferans, forepart of *Sclerolinum* and anterior trunk of female *O. priapus* [9,22,27,28]. Although the ciliary field in frenulates and both vestimentiferans and *Sclerolinum* lies in different regions, in all cases it originates from the larval neurothroch. In developing larvae of frenulate *Siboglinum fiordicum* the anterior part of neurotroch extended to the future forepart, whereas posterior part of neurotroch extended to the future trunk. In S. *fiordicum* only posterior part of neurotroch remains in the in trunk of adults [56,57]. Whereas in adult vestimentiferans it is in vestimentum corresponded to the frenulate forepart [9,27,58]. We assume that in adult frenulates and vestimentiferans different parts of the neurotroch remains, possibly due to different life modes of the larvae. Vestimentiferan larvae swim long time in the water, whereas in frenulates it settles and simultaneously goes through metamorphosis.

Perikarya do not form accumulations along the most length of the ventral nerve cord, i.e. in forepart/vestimentum and trunk, but their number increases in the region of annular chaetae, as in frenulate *Lamellisabella zachsi* [19,20] and in short opisthosomal segments of frenulate *Siboglinum fiordicum* perikarya form ganglia [21,22]. In contrast to vestimentiferans’ anchoring opisthosome, the frenulates’ opisthosome is designed to protrude out of the posterior tube opening and dig into the sediment [22]. Due to the high mobility, in the frenulate opisthosome the nerve cords form three strands with pair of ganglia in each segment in *Siboglinum fiordicum* [21,59].

Giant axons in vestimentiferans *Ridgeia, Riftia, Oasisia* [10,11,13] were found to extend up to the posterior end of the the trunk. In large frenulates like *Spirobrachia* and *Lamelisabella* there is a pair of giant axons extended from the giant unipolar perikarya located in the brain [20,22]. In small frenulates like *Nereilinum* there is only one giant axon, and it goes only along one side of the ventral ciliary field [22]. In frenulates the giant axons extend only untill the girdle of hook-shaped chaetae located approximately in the middle of the trunk, whereas in the vestimentiferans untill the end of the trunk. Giant axons provide a rapid contraction of the longitudinal musculature, serving as so-called “flight response” - in the frenulates and vestimentiferans it is the retraction of the body deep into the tube at the moment of danger (i.e. claws of crabs *Bythograea*). Frenulates anchored to the wall of the tube with means of girdle chaetae, and the vestimentiferans - the chaetae of opisthosome. That is why the giant axons reach only girdle in frenulates, and in the vestimentiferans - to the opistosome. There are no giant axons in *Osedax* and *Sclerolinum*.

Thus, the nerve cord in siboglinids is arranged in the similar way. In the anterior part of the body the paired strands of the nerve cord associated with the ventral ciliary field. In all groups, the nerve cord lies entirely within the epidermis and contains giant axons. The ventral nerve cord is not ganglionated for the most part of its length. The difference in the nervous systems is that the frenulates have a ganglionization in the opistosome, which probably indicates its greater mobility.

### Annelid ventral nerve cord in siboglinids

Siboglinids have intraepidermal paired medullary ventral nerve cord containing the giant axons (except *Sclerolinum* and *Osedax*) and associated with the ventral ciliary field. What features siboglinids share with the possible sister group of annelids?

Intraepidermal nervous system is also known in species of Opheliidae, Spionidae, Syllidae, Maldaniidae, Cossuridae, Polygordiidae, Protodrillidae etc as well as basal radiation Chaetopteridae, Magelonidae and Oweniidae [47,60–62]. Also, meiobenthic forms like Polygordiidae, Protodrilidae, Dinophiilidae have intraepidermal nervous system. So far, it is hard to tell the functional advantages of the intraepidermal nervous system or evolutionary aspects of it. Perhaps it is simply common among the sessile or meiobenthic forms.

Paired nerves in most annelids are found at the larval stages of Errantia, Sedentaria and their sister clade [48,52,63–69], whereas in adult annelids the nerve cord is organized in surprising range of levels: eighter single, paired, trineural, or pentaneural [47,60,70,71]. Based on presence of the paired nerve cords in the hypothetical sister clades Cirratulida and Sabellida [39,72] and paired organization of the nerve cord in siboglinids, we can conclude that the paired nerve strands within the ventral nerve cord might be ancestral feature for siboglinids (Fig 12).

Lack of ganglia in medullary nerve cord in long vestimentum/forepart and trunk segments of vestimentiferans and frenulates, and their presence in each segment of mobile frenulate opisthosome is unusual for the most annelids exhibiting the uniform structure of the nerve cord along worm body as eighter medullar, or ganglionated one [47,60,70,71]. Non-uniform ventral nerve cord is known in oweniids: nerve cord exchibits medullary state in elongated anterior segments and ganglionated-like state in short posterior segments [44,52,73]. We assume in siboglinids medullary state of nerve cords in elongated segments is due to regular innervation of the structures in the segments which is convergent to the state of oweniid nerve cord.

The pattern of the segmental neurite bundles in vestimentiferans is similar to what we know in oweniids [44,52,73]: numerous and anastomosing in long segments and condensed single bundle in short opisthosomal segments. This pattern could be a reflection of the elongation of the segments.

Giant axons and giant perikarya are common among annelids, mainly in large forms [71,74]. Common feature of most annelids to have multicellular or unicellular giant fibres extending from the giant somata usually lying in subesophageal ganglia and/ or other segmental ganglia. In vestimentiferans it is known a pair of giant perikarya, whereas in *Riftia* we detected at least four pairs of somata lying in the area of supraesophageal ganglion. Among annelids only in sabellids, like large *Myxicola infundibulum* and *Sabella pavonina*, the giant perikarya lie in supraesophageal ganglion [74]. So, the vestimentiferans share with sabellids the similar position of the giant perikarya in the supraesophageal ganglion.

Vestimentiferans together with the rest siboglinids have the ventral ciliary field bordered by a pair of strands of the nerve cord. The ciliary field is not common among sexually matured annelids. The structure is known in progenetic *Dinophilus gyrociliatus* [75,76] and used for gliding. There is an observation that tiny frenulate *Nereilinum murmanicum* uses the ciliary field to glide vertically along its tube [23]. Other functions of the ventral ciliary fields in siboglinids remains theoretical [5]. Thus, it is the paired sturucture of the ventral nerve cord that siboglinids share with the possible annelid sister groups (Fig 12).

### Brain ogranization in vestimentiferans

The differences in brain structure of vestimentiferan species are mainly in the shape of their brains and the presence/absence of cuticule structures [6,7,10–12,15,54]. *Riftia pachyptila*’s brain has heart-like shape at the transverse section with significatly developed dorso-lateral lobes (Fig 5, 8A-D). Brain of *Ridgeia piscesae* has triangular shape at transverse section with wide ventral side [10]. Brain of *Lamellibrachia luymesi* has oval transverse shape [6]. These two latter vestimentiferan species have less developed dorso-lateral lobes in comparison to *Riftia* (S5 Fig). *Rifia* is known to possess 340 tentacles per lamellae and 335 lamellae on each side of the obturaculum whereas 70 lamellae in *Escarpia* is the maximum lamellae number among the rest vestimentiferans [7,77,78]. This could be the explanation of the presence of the enlarged dorso-lateral lobes in *Riftia’s* brain. Notably, inspite of the brain shape differences, tentacle nerves originate from the same dorso-lateral areas of the brain neuropile in *Riftia* and all other vestimentiferans.

Cuticle schield protects ventral side of the brain that has a direct contact with the tube or ambient environment in all studied vestimentiferans as well as in *Riftia* (Fig 10G) [6,10,11,15]. The dorsal and frontal sides of the brain are covered by tentacles and obturacules (Figs 4, 5). Additionaly, brain can be penetrated by cuticle shifts and plates extending from the cuticle of tentacle lamellae, as in *L. luymesi*, *R. piscesae*, *O. alvinae*, but not in *Riftia* [6,11,15].

### Annelid brain in vestimentiferans

The juvenile vestimentiferans preserve the gut rudiment what assist to make homologization of the brain parts of the gutless siboglinids with the supra- and subesophageal ganglia of typical annelids.

The brain of the vestimentiferans lies completely in the epidermis at the anteriormost part of the vestimentum. It is the large and dense mass of the neuropile which looks like single entity, non-subdivided into the supraesophageal and subesophageal ganglia, as in most annelids [60]. Following the idea suggested by Jones and Gardiner [8] we assume that to the part of the brain of the vestimentiferans lying dorsally to the enteral coelomic channel can be homologized with the supraesophageal ganglion (Figs 4, 8 A D, 11), whereas the part of the brain lying ventrally to the enteral coelomic channel - with the subesophageal ganglion (Figs 4, 8 E-H, 11).

**Fig 11.**
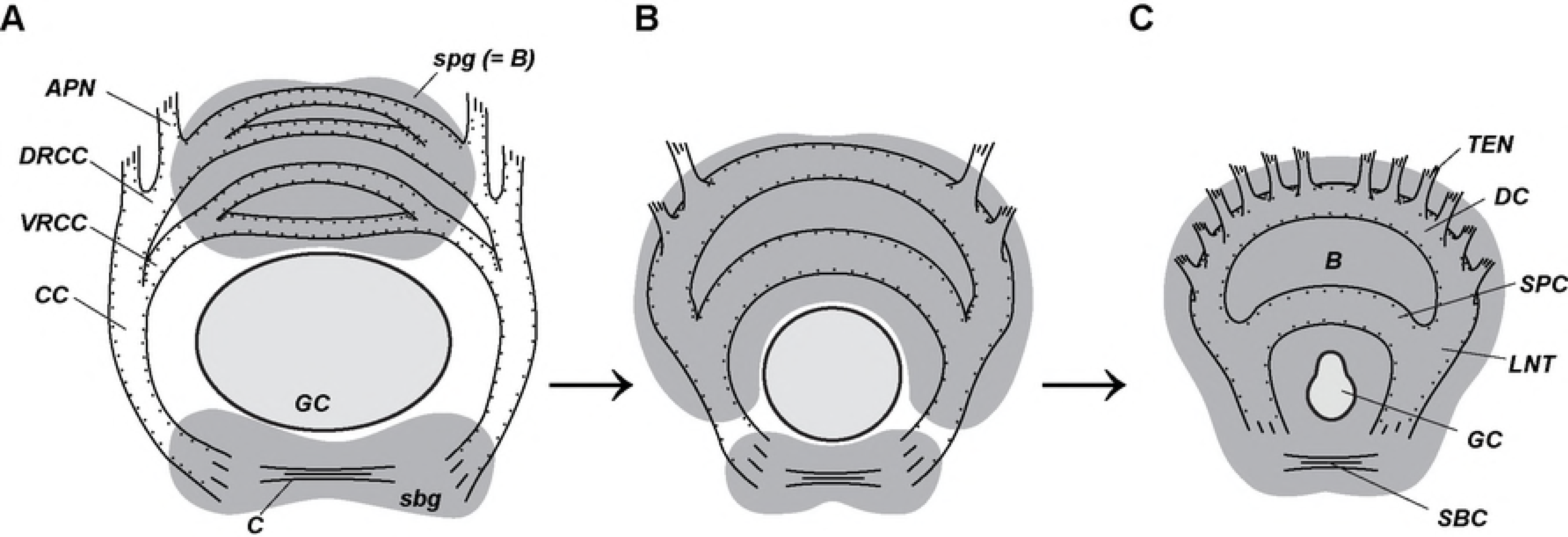
Hypothetical vestimentiferan brain origin. A - supra- and subesophageal ganglia in annelids (after [47]). B - hypothetical transitional state. C - vestimentiferan brain. *APN* – neurite bundles of palps, *B* – brain, *C* – commissure of *sbg, CC* – circumesophageal connectives, *DC* – dorsal commissure, *DRCC* – dorsal (posterior) root of the CC, *GC* – enteral coelom, *LNT* – longitudinal nerve tracts projecting from the ventral nerve cord into the brain, *sbg* – subesophageal ganglion, *SBC* – subenteral commissure, *spg* – supraesophageal ganglion (which is brain in annelids), *SPC* – supraenteral commissure, *VRCC* – ventral (anterior) root of CC, *TEN* – neurite bundles of tentacles (palps).

Longitudinal nerve tracts (*LNT*) lie ventrally in the posterior brain, while in the anterior brain tracts run symmetrically right and left to the enteral coelomic channel and connect each other by transverse commissures in the dorsal part of the anteriormost part of the brain. Thus, LNT can be homologized with the circumesophageal connectives of annelids (Figs 9, 11). In annelids the circumesophageal connectives enter the brain and bifurcate into ventral and dorsal roots [43,79–83]. Each root connects by a pair of dorsal and ventral commissures. Thus, in the annelid supraesophageal ganglion, there are two pairs of transverse commissures: a dorsal pair and ventral one [43,47,81,83,84]. In vestimentiferans’ dorsal brain part two transverse commissures can be distinguished: the dorsal commissure consisting of couple of transverse bundles, and the supraenteral commissure. Both commissures connect the nerve bundles of the longitudinal nerve tracts. We suppose that the dorsal and supraenteral commissures of the brain of vestimentiferans can be homologized with dorsal and ventral pairs of the commissures of the supraesophageal ganglion of typical annelids (Fig 11).

Posteriorly, longitudinal nerve tracts pass through the ventral part of the brain and come into the ventral nerve cord as the circumesophageal connectives in the annelid brain and continue as longitudinal connectives of the ventral nerve cord [84].

The innervation of numerous tentacles of *R. pachyptila* occurs from the neuropile of the lateral brain lobes (NE) containing radial tentacle neurite bundles (Figs 7, 8 A-D, 11C). Neuropiles of the lateral lobes adjoin the longitudinal nerve tracts which are possible homologues to the circumesophageal connectives (Fig 11). In annelids, the most part of the peristomial palps are innervated from the circumesophageal connectives [43,47]. Previously, tentacles of vestimentiferans were homologised with palps of polychates [31], although based on differences in the external and internal structures (lack of ciliated grooves, absence of longitudinal support rods and the presence of the afferent and efferent blood vessels inside each tentacle) this homology was considered as doubtful [58]. Our data on the innervation of the tentacles of *Riftia* proves the annelid palps hypothesis of the vestimentiferan tentacles (Fig 11). But in the vestimentiferans (especially *Riftia*) the parts of longitudinal nerve tracts and neuropile of the lateral brain lobes are incomparably larger then corresponding neural structures in annelids, because the tentacle apparatus of vestimentiferans is significantly developed. The similar correlation between sizes of the tentacle crowns and brains are clearly seen in oweniids and sabellids. The brain of oweniids with simple gill tentacles is just a transverse commissure passing in the epidermis dorsal to the digestive tract [44,73], whereas in sabellids with the large complecated tentacle crown serving for food collection the brain consists of main four transverse commisures and many other additional neural structures [43].

To summarize, the vestimentiferan brain shows similarity to the annelid brain organization if we accept the idea of Jones and Gardiner [8] that the brain is a result of the union of the supra- and subesophageal ganglia. In the dorsal part of the vestimentiferan brain (=supraesophageal ganglion) we found homologues of the dorsal and ventral pairs of the transverse commissures. The annelid brain shows remarkable variety of the organization [46,82,85,86]. Our comparative anatomical approache shows that the structure of the vestimentiferan brain and nervous system does not go beyond this diversity of the brain and the nervous system of annelids.

Selivanova et al. [87] conducted a single immunoreactive study on brains of vestimentiferan *Ridgeia piscesae* and identified 60 FMRFamide-immunoreactive neurons in posterior brain and 24 neurons in ventral part of the brain and single FMRF-amide IR-like processes in the medial zone of the brain neuropile. These specific FMRFamide IR-like correspond to the following components of the brain of *Riftia pachyptila:* posterior median perikarya aggregations (*pmp*, Figs 7; 8E-H; S6F Fig), ventrolateral lobe and posterior part of ventral lobe of tripartite ventral aggregation of perikarya (*lvtp, vvtp*, Figs 5; 6; 7; 8E-H; S6 A-D Figs) and vertical median bundles (*VAN, VPN*, Figs 6; S10 Fig). The effect of FMRFamide mediator are shown to support the heart pulsation, and tone of the esophagus and body walls in *Nereis virens* and *Sabellastarte magnifica* [88,89]. In molluscs, FMRF-amide mediator is known to excite and inhibit heartbeat [90], while in insects it controls heart function, somatic musculature, crop and salivary glands [91]. Indeed, FMRFamide IR-elements in the brain of *Riftia* are close to coelomic channels containing the rudimentary gut and obturacule blood vessels (*pmp, VAN, VPN*, Figs 6, 7) and to the ventral wall of the body (*lvtp, vvtp*, Figs 5; 6; 7). Perhaps the mentioned brain components of the vestimentiferan brain are related to the functioning of the heart, gut and musculature of the body wall.

### Evolutionary aspects of siboglinid brain

Vestimentiferans of genera *Riftia, Ridgeia, Oasisia, Lamellibrachia* and *Osedax* are the only siboglinids so far, whose brains were studied in detail [5,9,11,12,15,26,27,87] and present study]. Their brains compeletely lying in the ventral body epidermis and so far in spite of the fact that *Osedax* does not have gut rudiment as *Riftia*, it is still possible to find similar structures helping us to homologize parts of brains among siboglinids.

First, homologization of dorsalmost commissures in brain of *Riftia* (*DC*) and anterior commissure in *Osedax* (*ACBR*). They could be homologous to each other based on anteriormost position in the brains in both siboglinids as well as based on neurites originating from them. Various vertical neurite bundles originating from these dorsalmost commissures in vestimentimentiferans and *Osedax* (*DC* in *Riftia* and *ACBR* in *Osedax*) and innervate anterior structures: 1) obturacule neurites in vestimentiferans (*OBN* in *Riftia*) and antero-dorsal nerves or anterior nerve net in *Osedax (ADN* and *ANN*, Fig 12; see Fig 2 in Worsaae et al., 2016), 2) palp neurites in *Riftia* (*TEN*) and *Osedax* (*PN*). Second, homologization of palp neurites in *Riftia* (*TEN*) and *Osedax* (*PN*) based on the similar origin of lateral parts of the anterior most commissures in *Riftia* (*DC*) and *Osedax* (*ACBR*, Fig 12).

**Fig 12.**
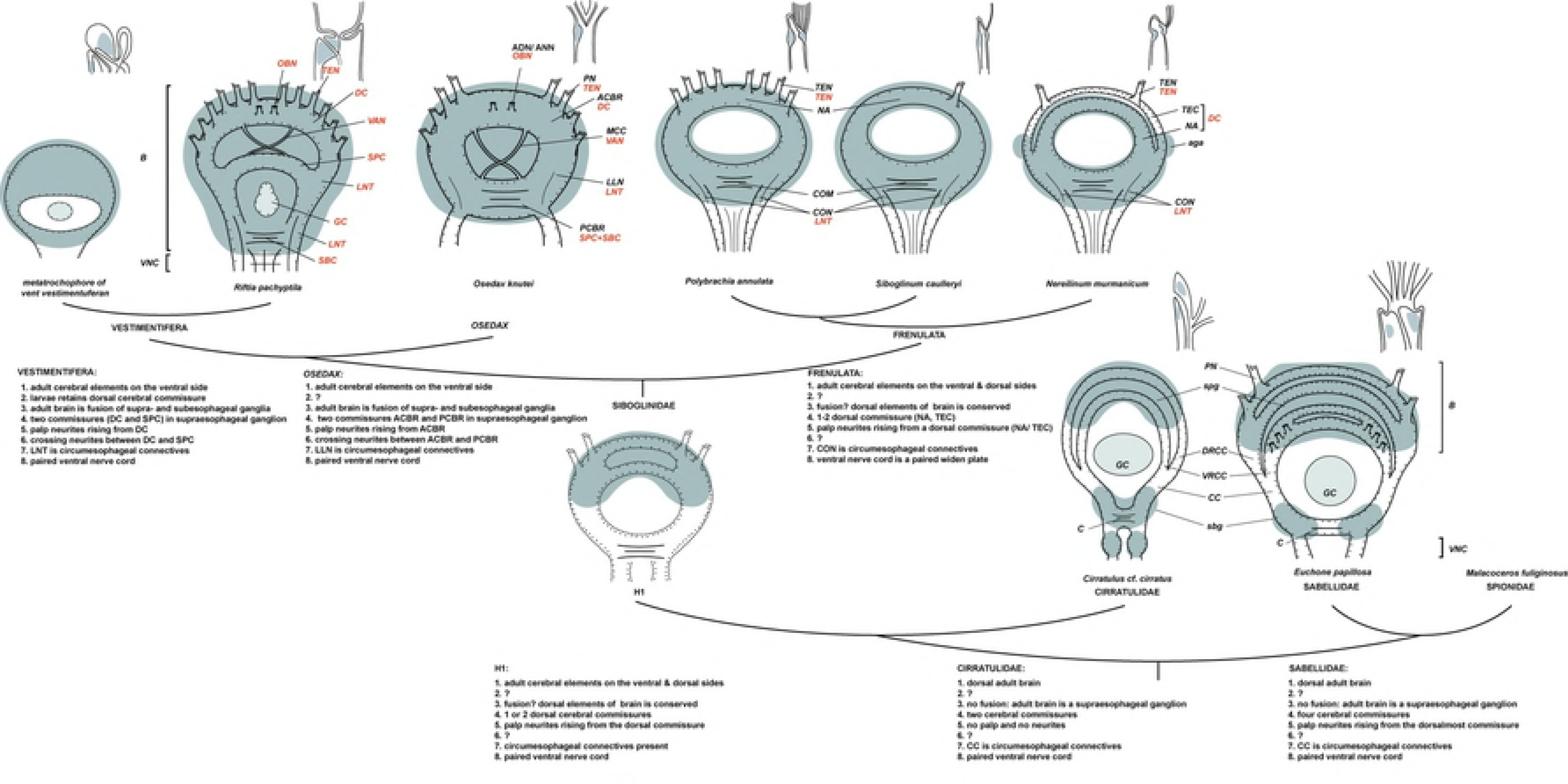
Reconstruction of hypothesized neural ancestor traits of siboglinid central nervous system. Relation among the siboglinid groups from a combination of recent phylogenetic trees: clade of frenulates resolved based on cladistic analysis [18], interrelationship of siboglinid clades based on phylogenetic and phylogenomic data [17,92]; annelid outgroups based on phylogenomic data [39]. Neural characters for homologization of cerebral elements of siboglinids and annelid sister groups are listed (1-8). Neural diagrams include sagittal views (upper row) and dorsal views (lower row) with disposition of the cerebral elements in the anteriormost end of the worms. Perikarya shown in gray blue. Anterior ends at the top. Dashed lines show hypothesized brain boundary in the siboglinid ancestor. Cerebral elements drawn based on larvae of vestimentiferan *Riftia pachyptila* [14], adult of vestimentiferan *Riftia pachyptila* (this study); *Osedax knutei (=O*. “nudepalp E”, [27,93]); frenulates *Polybrachia annulata*, and *Siboglinum caulleryi* [19,20], *Nereilinum murmanicum* [23], cirratulid *Cirratulus cf. cirratus* [50], sabellid *Euchone papillosa* [43]. *Sclerolinum*’s cerebral elements of the ventral brain are not shown. H1 is a combination of the hypothesized ancestral siboglinid states. *ACBR* - anterior commissure of the brain, *ADN* – antero-dorsal nerve, *aga* – ganglion-like aggregation at base of *TEC, ANN* - anterior nerve net, *B* – brain, *C* – commissure in *sbg, CC* – circumesophageal connectives, *COM* – commissure, *CON* – connective, *DC* – dorsal commissure, *DRCC* – dorsal (posterior) root of the CC, *GC* – enteral coelom, *LLN* – lateral longitudinal neurite bundles in the brain, *LNT* – longitudinal nerve tracts projecting from the *VNC* into the brain, *MCC* – middle cross commissure, *NA* – nerve ring (after [20]) or brain ring (after [23]), *OBN* – obturacular neurites, *PCBR* – posterior commissure of the brain, *PN* – palp nerve, *TEC* – tentacular commussure, *TEN* – neurite bundles of tentacles (palps), *SBC* – subenteral commissure, *sbg* – subesophageal ganglion, *SPC* – supraenteral commissure, *spg* – supraesophageal ganglion, *VAN* – anterior vertical median bundles, *VNC* – ventral nerve cord, VRCC – ventral (anterior) root of CC.

Third, homologization of the supraenteral commissure in *Riftia* (*SPC*) and the anterior part of posterior commissure in *Osedax (PCBR*, Fig 12). In the brain of *Riftia* (present study) and *Osedax* [27] there are crossing neurites: in the median areas of the brains in *Riftia* (*VAN*) and *Osedax* (*MCC*). Based on the presence and position of crossing neurites, we consider these commissures which are connected by crossing neurites homologous to each other. This means that the thick posterior commissure in *Osedax* (*PCBR*) is homologous to the union of supra- and subenteral commissures in vestimentiferan brain (Fig 12). So, if in *Osedax* gut rudiment has been remained, it would pass throught the *PCBR*.

Forth, based on listed above homologozations we consider the longitudinal nerve tracks in *Riftia* (*LNT*) are possibly homologous to lateral longitudinal bundles in *Oseadx* (*LLN*), and both are homologous to the circumesophageal connectives in the brain of annelids (Fig 12).

Organization of the brain of frenulates, sister group to all other siboglinids [2,92] are important for the analyisis of the ancestral state of the siboglinid brain. Although their brains were studied in less details, but it is known in the dorsal epidermis in *Polybrachia annulata, Siboglinum caulleryi* there is a dorsal commissure [19,20], and in *Nereilinum murmanicum, S. modestum* and S. *subligatum* there are two dorsal commissures [23]. These commissures in frenulates give rise the neurite bundles to anterior appendages. We consider that these dorsal commissures (*NR, TEC*) bearing anterior appendages’ nerves in frenulates (Fig 12) could be compared with the anterior commissures in ventral brain of vestimentiferans and *Osedax* (*DC* and *ACBR*, respectively) bearing nerves to palps. Moreover, we assume that according to innervation pattern, the anterior appendages in frenulates are also annelid palps.

Brain of *Sclerolinum* is very simple structure lying completely on the ventral side and having two layeres: apical perikarya and basal neuropile [28]. Due to the described simpilicity of the brain structure we do not homologize it with other siboglinids and annelids.

Intriguing question is what the ancestral state of the brain of siboglinid was and how it was evolved? Besides the frenulates, vestimentiferan metatrochophores have the commissures and perikarya in the dorsal epidermis [14]. And sister clades to Siboglinidae, Cirratulidae and Sabellidae (according to [39]), have well developed supraesophageal brain on the dorsal side of the body [43,50]. It is reasonable to hypothesize that the presence of the commissure, lying dorsally to the gut and giving rise neurite bundles to palps might be the ancestral state of the siboglinid brain (Fig 12).

## Conclusions

Our microscopical study and 3D reconstruction of the central nervous system of the giant vestimentiferan tubeworm *Riftia pachyptila* allowed to reveal the structure of the brain and ventral nerve cord.

Brain in adult vestimentiferans is a product of fusion of supraesophageal and subesophageal ganglia. In the part homologized with the supraesophageal ganglion there are two commissures: double dorsal commissure and single supraenteral commissures. In the subesophageal part there is the subenteral commissure. Based on the innervation of the anterior appendages from the longitudinal nerve tracts which are homologous to the circumesophageal connectives, tentacles of vestimentiferans are annelid palps. The innervation of the obturacles is different and will be discussed in the following publication. The ventral nerve cord of vestimentiferans is intraepidermal, paired, associated with the ciliary field, not ganglionated for the most part of its length. The latter is the feature of the elongation of the segments. In *Riftia* there is one giant axon in the ventral nerve cord in trunk which is the product of fusion of several axons. The giant axon extends from at least four giant perikarya in Riftia. Suprisingly, the giant perikarya lie in the supraesophageal brain part of *Riftia*, like in sabellid annelids.

The central nervous system of vestimentiferans and other siboglinids are arranged in the same way: anterior ventral brain and paired ventral nerve cord. All siboglinids share the same features of the ventral nerve cord: intraepidermal paired nerve cord associated with the ventral ciliary field (Fig 12). The comparative analysis of brain structures of the siboglinids suggests that the dorsal commissure bearing palp nerves is common for all siboglinids: it can be found in dorsal epidermis of frenulates and vestimentiferan larvae, in supraesophegeal part of vestimentiferan brains and possibly it is anterior commissure found in *Osedax* (Fig 12). The difference in the nervous systems is that the vestimentiferans have a large and significatly differentiated brain which is reflection of the high development of the palp apparatus. *Osedax*, frenulates and *Sclerolinum* have less developed brain. Frenulates and *Sclerolinum* have good ganglionization in the opisthosome, which probably indicates its high mobility.

The comparative anatomical analysis of the neural structures of the siboglinids and sister annelid clades lead us to hypothesize that the presence of the commissure, lying dorsally to the gut and giving rise neurite bundles to palps might be the ancestral state of the siboglinid brain.

## Acknowledgments

We thank the chief scientists of Laboratory of Ocean Benthic Fauna and crews of RVs and ROVs of Shirshov Institute of Oceanology of the Russian Academy of Science for assistance and their continuous help in collecting the material.

## Supporting Information

### S1 Figure. Anteriormost brain of *Riftia*

Histological cross section of the 79 mm long male. Level of the section shown at the diagram, right lower coner. *amp* – anterior median aggregation of perikarya, *CU* – cuticle, *CUP* – cuticle schild, *DC* – dorsal commissure, *DLN* – dorsal area of the longitudinal bundles, *dop* – dorsal aggregation of perikarya, *EP* – epidermis, *GC* – enteral coelom, *LNT* – longitudinal nerve tracts projecting from the ventral nerve cord into the brain, *LR* – undifferential tentacle lamellae, *NE* – neuropile of the lateral brain lobes, *nep* – peripheral perikarya of the lateral brain lobes, *OBC* – obturacular coelom, *OBL* – obturacular lobes, *OBN* – obturacular neurite bundles, *OBV* – obturacular blood vessels, *SLN* – supraenteral longitudinal neurite bundles, *TE* – free tentacles, *TEN* – neurite bundles of tentacles (palps), *vtp* - tripartite ventral aggregation of perikarya.

### S2 Figure. Anterior and middle brain organization of *Riftia*

A-B - histological cross sections of the 79 mm long male. Level of each section shown at the diagram, right lower coner. *amp* – anterior median aggregation of perikarya, *CUP* – cuticle schild, *DC* – dorsal commissure, *DLN* – dorsal area of the longitudinal neurite bundles, *dop* – dorsal aggregation of perikarya, *EP* – epidermis, *GC* – enteral coelom, *LNT* – longitudinal nerve tracts projecting from the ventral nerve cord into the brain, *LR* – undifferential tentacle lamellae, *lvtp* – ventrolateral perikarya of the *vtp, NE* – neuropile of the lateral brain lobes, *nep* – peripheral perikarya of the lateral brain lobes, *OBL* – obturacular lobes, *OBC* – obturacular coelom, *OBN* – obturacular neurite bundles, *OBV* – obturacular blood vessels, *pl* – large perikarya, *ps* – small perikarya, *SLN* – supraenteral longitudinal neurite bundles, *TE* – free tentacles, *TEN* – neurite bundles of tentacles (palps), *VSN* – vertical supraenteral neurite bundles, *vtp* - tripartite ventral aggregation of perikarya, *vvtp* - ventral perikarya of the *vtp, XXL* – pair of prominent bundles of large longitudinal neurites (part of *LNT*).

### S3 Figure. Posteriormost brain organization of *Riftia*

A-B - histological cross sections of the 79 mm long male. Level of the section shown at the diagram, right lower coner. *CUP* – cuticle schild, *DC* – dorsal commissure, *DLN* – dorsal area of the longitudinal neurite bundles, *dop* – dorsal aggregation of perikarya, *EP* – epidermis, *GA* – giant axons, *GC* – enteral coelom, *LNC* – lateral connectives, *LNT* – longitudinal nerve tracts projecting from the ventral nerve cord into the brain, *LR* – undifferential tentacle lamellae, *lvtp* – ventrolateral perikarya of the *vtp, NE* – neuropile of the lateral brain lobes, *nep* – peripheral perikarya of the lateral brain lobes, *OBC* – obturacular coelom, *OBL* – obturacular lobes, *OBN* – obturacular neurites, *OBV* – obturacular blood vessels, *pl* – large perikarya, *pmp* – posterior median perikarya aggregation, *ps* – small perikarya, *SBC* – subenteral commissure, *SPC* – supraenteral commissure, *SLN* – supraenteral longitudinal neurite bundles, *TEN* – neurite bundles of tentacles (palps), *VPN-* posterior vertical median bundles, *vvtp* - ventral perikarya of the *vtp, VSN* – vertical supraenteral neurite bundles.

### S4 Figure. Intraepidermal position of the brain

A, B - schemes of sagittal and cross sections at levels (1-3) shown in (A). *B* – brain, *CU* – cuticle, *CUP* – cuticle schild, *ECM* – extracellular matrix, *EP* – epidermis, *OBL* – obturacular lobes, *VE* – vestimental process, *VNC* – ventral nerve cord.

### S5 Figure. Neural elements in the bases of the undifferential tentacle lamellae

A - scheme of neural elements of the undifferential tentacle lamellae: perikarya and neurite bundles. B-D - tentacle lamellae bases on the dorsal, lateral and ventrolateral sides of the brain surface, respectively. *ECM* – extracellular matrix, *EP* – epidermis, *IEP* – epidermis of the internal lamellae wall, *OEP* – epidermis of the external lamellae wall, *LR* – undifferential tentacle lamellae, *NE* – neuropile of the lateral brain lobes, *nep* – peripheral perikarya of the lateral brain lobes, *NB* – neurite bunldes, *OB* – obturaculum, *pl* – large perikarya, *ps* – small perikarya, *TEN* – neurite bundles of tentacles (palps).

### S6 Figure. Coelomic channels running through the brain

3D models of *Riftia* brain. A, C, E - peripheric perikarya of the lateral brain lobes (*nep*) are on the dorsal side of the brain (purple). B, D, F - tripartite aggregation of perikarya (*vtp*) is on the ventral side and under the obturacular and enteral coeloms (blue). View sides shown at the right lower corners of each images. Cube side is 255 μm. Dashed lines point neural elements under transparent structures. *GC* – enteral coelom, *lvtp* – ventrolateral perikarya of the *vtp, nep* – peripheral perikarya of the lateral brain lobes, *OBC* – obturacular coelom, *pmp* – posterior median perikarya aggregation, *vtp* - tripartite ventral aggregation of perikarya, *vvtp* - ventral perikarya of the *vtp*.

### S7 Figure. Innervation of neuropile of the lateral brain lobes

3D models of *Riftia* brain. A-C, A’-C’ - neuropile of the lateral brain lobes (*NE*) associated with the longitudinal nerve tracts (*LNT*). D-H - longitudinal nerve tracts projecting from the ventral nerve cord into the brain (*LNT*) and giving rise the prominent bundles of large longitudinal neurites (*XXL*). I-J - peripheric perikarya (*nep*) and neuropile of the lateral brain lobes (*NE*). View sides shown at the right lower corners of each images. Cube side is 255 μm. Dashed lines point neural elements under transparent structures. *DC* – dorsal commissure, *DLN* – dorsal area of the longitudinal bundles, *GA* – giant axons, *GC* – enteral coelom, *LNC* – lateral connectives, *LNT* – longitudinal nerve tracts projecting from the ventral nerve cord into the brain, *NE* – neuropile of the lateral brain lobes, *nep* – peripheral perikarya of the lateral brain lobes, *SBC* – subenteral commissure, *SPC* – supraenteral commissure, *vtp* - tripartite ventral aggregation of perikarya, *XXL* – pair of prominent bundles of large longitudinal nerve tracts (part of *LNT*).

### S8 Figure. 3D-models of anterior neural elements of *Riftia* brain

3D models of *Riftia* brain. A-C – overviews of supraenteral longitudinal neurite bundles (*SLN*) extending from the anterior median perikarya aggregation (*amp*); D-E - anterior median perikarya aggregation in association with the main cerebral elements: ventral tripartite aggragation (*vtp*), dorsal commissure (*DC*) and supraenteral commissure (*SPC*); F-G - anterior median perikarya aggregation (*amp*) and dorsal commissure (*DC*) in association with the obturacular channels (*OBC*). View sides shown at the right lower corners of each images. Cube side is 255 μm. Dashed lines point neural elements under transparent structures. *amp* – anterior median aggregation of perikarya, *DC* – dorsal commissure, *GA* – giant axons, *GC* – enteral coelom, *LNC* – lateral connectives, *OBC* – obturacular coelom, *SBC* – subenteral commissure, *SPC* – supraenteral commissure, *SLN* – supraenteral longitudinal neurite bundles, *VSN* – vertical supraenteral neurite bundles, *vtp* - tripartite ventral aggregation of perikarya.

### S9 Figure. Obturacular innervation and giant neurons in *Riftia* brain

3D models of *Riftia* brain. A-C – disposition of obturacular neurite bundles (*OBN*) and dorsal longitudinal bundles (*DLN*), D-F – origin of obturacular neurite bundles (*OBN*) from dorsal commissure, and neuropile of the lateral brain lobes (*NE*) from longitudinal nerve tracts (*LNT*). G-J – giant axons (*GA*) and position of giant perikarya (*gap*); K-L – position of giant neurons between the coelomic channels (*OBC, GC*). View sides shown at the right lower corners of each images. Cube side is 255 μm. Dashed lines point neural elements under transparent structures. *DC* – dorsal commissure, *DLN* – dorsal area of the longitudinal bundles, *GA* – giant axons, *GC* – enteral coelom, *gap* – giant perikarya, *LNT* – longitudinal nerve tracts projecting from the *VNC* into the brain, *NE* – neuropile of the lateral brain lobes, *nep* – peripheral perikarya of the lateral brain lobes, *OBC* – obturacular coelom, *OBN* – obturacular neurites, *SPC* – supraenteral commissure, *XXL* – pair of prominent bundles of large longitudinal nerve tracts (part of *LNT*).

### S10 Figure. Vertical midbrain neurite bundles

3D models of *Riftia* brain. A-D - anterior (*VAN*) and posterior (*VPN*) vertical median bundles in between other midbrain structures, E-G - giant axons (*GA*) running inside the crossing anterior median bundles (*VAN*). View sides shown at the right lower corners of each images. Cube side is 255 μm. Dashed lines point neural elements under transparent structures. *DC* – dorsal commissure, *GA* – giant axons, *GC* – enteral coelom, *gap* – giant perikarya, *LNC* – lateral connectives, *OBC* – obturacular coelom, *SBC* – subenteral commissure, *SLN* – supraenteral longitudinal neurite bundles, *SPC* – supraenteral commissure, *VAN* – anterior vertical median bundles, *VPN* - posterior vertical median bundles.

